# A machine learning framework for evolutionary model classification

**DOI:** 10.64898/2026.11.20.768158

**Authors:** Menna Jones

## Abstract

Machine learning algorithms are increasingly used across scientific disciplines, but remain absent from several areas of comparative biology. One example is evolutionary model selection, which typically relies on likelihood-based methods to identify the model among a candidate set that best explains trait evolution across a phylogeny. This approach requires all candidate models to have a tractable likelihood function yet this may not the case for complex models. Alternative likelihood-free inference methods are available but their application is limited in scalability and flexibility. Both classes of methods struggle with discriminating models that yield statistically similar patterns in trait data. Here, I introduce a machine learning framework to perform evolutionary model classification from comparative data. Using simulated datasets, I demonstrate that the approach improves model discriminability relative to conventional likelihood-based model fitting but does not overcome model non-identifiability. I then revisit an analysis of early burst signals in comparative data to assess how a machine learning approach alters the patterns of early burst support obtained through likelihood-based inference. Finally, I discuss future extensions of this framework and their potential to improve both our representation and understanding of evolutionary process-to-pattern mapping.

## Introduction

Quantitative models of evolutionary processes are central to modern comparative biology, from inference of diversification dynamics (Nee et al., 1994; Morlon et al., 2010, 2011; Rabosky, 2014; Maddison et al., 2007; FitzJohn et al., 2009; FitzJohn, 2010; Louca and Pennell, 2020; Stadler, 2010) to tempo and mode in trait evolution (Felsenstein, 1973; Hansen, 1997; Butler and King, 2004; O’Meara et al., 2006; Boucher and Démery, 2016; Clavel and Morlon, 2017). Evaluating which model best explains an observed set of evolutionary patterns is essential for linking statistical inference to evolutionary theory and is usually achieved by specifying and optimizing a likelihood function (Edwards, 1972) for each model. The resulting parameter estimates are those that maximize the likelihood of observing the data under the model, while the maximized likelihoods are measures of support that can subsequently be used to perform model selection via likelihood ratio tests, or else transformed into information criteria, such as the Akaike (AIC, Wagenmakers and Farrell, 2004) or Bayesian information criterion (BIC, Schwarz, 1978) to mitigate bias favoring highly-parameterized models and aid in evaluation of relative model support.

Likelihood-based approaches are limited by the need for a tractable probability density function (PDF) (Diggle and Gratton, 1984) for each evaluated model. This requirement is circumvented by likelihood-free statistical inference techniques. For example, Approximate Bayesian Computation algorithms (ABC, (Rubin, 1984; Beaumont et al., 2002)) approximate parameter posterior distributions through rejection-based MCMC sampling of parameters that minimize the distance between simulated and observed data according to a chosen distance metric (Lintusaari et al., 2017). In evolutionary biology, ABC has seen some use in studies of diversification dynamics and trait evolution (Slater et al., 2012; Rabosky, 2009; Xie et al., 2026; Jhwueng and Chang, 2025) but faces non-trivial limitations in computational efficiency and scalability to large datasets (Green et al., 2015). These limitations are becoming increasingly relevant with the development of larger and more complex datasets across scientific disciplines, including phylogenetic comparative biology (Jarvis et al., 2014; Fleming et al., 2023; Steenwyk et al., 2023), and motivate a shift toward a more flexible, algorithmically-focused modeling framework (Breiman, 2001).

Machine learning (ML) algorithms are the subset of artificial intelligence algorithms that perform pattern recognition and classification tasks typically associated with human cognition (McCarthy and Hayes, 1981; Mitchell, 1997). Unlike conventional statistical approaches, ML algorithms capture nonlinear relationships and interactions among predictors without requiring an explicit likelihood function or known functional relationship between them. This flexibility makes ML algorithms well-suited to handle complex, high-dimensional datasets. Although classifier training can be computationally expensive, this cost is incurred only once, after which the classifier is broadly applicable across unseen classification scenarios without requiring additional computational effort. In evolutionary biology, ML is seen in contexts ranging from microfossil classification (see Yaqoob et al., 2025, and references therein), to three-dimensional morphometric analyses (Percival et al., 2019; Porto et al., 2021) and epidemiological phylodynamics (Voznica et al., 2022; Lambert et al., 2023). Bokma (2006) used an artificial neural network to estimate parameters of phenotypic evolution under Brownian motion (BM: Felsenstein, 1973; Harvey and Pagel, 1991) but did not extend to the task of model discrimination. More recently, Roa Lozano et al. (2026) introduced Evolutionary Discriminant Analysis (EvoDA) as the first explicit ML framework for evolutionary model classification and found it to outperform likelihood-based model-fitting approaches in the empirical context of fungal trait evolution across an 18-species phylogeny.

ML approaches may overcome several limitations of likelihood-based methods for modeling continuous trait evolution. First, likelihood-based methods are data-hungry: even simple models can be difficult to distinguish on small trees (Boettiger et al., 2012; Cooper et al., 2016b), and complex models may require larger datasets than most comparative biologists have (Beaulieu et al., 2012). Second, because BM is a special case of all alternative models, model classification becomes increasingly difficult as the magnitude of model parameters decreases and the process begins to look more like BM (Hansen, 1997; Cooper et al., 2016b; Slater and Pennell, 2014; Cressler et al., 2015). Finally, multiple evolutionary mechanisms can generate similar comparative signatures (Foote, 1996a; Revell et al., 2008). This many-to-one mapping from process to pattern can hinder model discriminability or produce erroneously strong support for a model that implies a process very different from the true generative one (Jones and Slater, 2026). Some models may even be non-identifiable when working with ultrametric trees. For example, for every value of the rate acceleration parameter *a* in the Accelerating Rates (AC) model there is a single peak OU model with the same likelihood when *α* = 2*a*, such that this models cannot be distinguished in the absence of fossil data (Uyeda et al., 2015).

In this study, I introduce a feed-forward multi-layer perceptron (MLP) neural network framework for classifying comparative data to their generating statistical model. I outline the statistical foundations of the classifier before describing the simulation, parameterization, and training procedures used to construct it. I then evaluate its predictive performance against conventional maximum likelihood (MaxL) model selection and explore individual predictor contributions using Bayesian Additive Regression Trees (BART). Finally, I revisit the study of Harmon et al. (2010b) to determine whether the prevalence of early burst dynamics in comparative data can be reassessed in a MLP classification framework.

## Materials & Methods

All analyses were conducted in R v. 4.4.1 (R Core Team, 2024). Model definition, training, and application were performed using functions of the torch package (Falbel, 2020) for R, which is adapted from the PyTorch architecture (Paszke et al., 2019).

To ensure that the MLP generalizes well across evolutionary scenarios and datasets, training data must cover a wide range tree topologies and data, generated with plausible parameter combinations. I generated 2 × 10^3^ birth-death phylogenies with pbtree() in the phytools package (Revell et al., 2024), drawing birth rates from *λ* ∼ U(0.5, 2) and death rates as *µ* = *λu* with *u* ∼ U(0.05, 0.5). Trees contained 20-1000 extant tips and were scaled to unit depth. Univariate traits were then simulated on each tree with fastBM() (Revell et al., 2024) under five models: (1) Brownian motion (BM Felsenstein, 1973), in which variance accumulates linearly at rate *σ*^2^; (2) a single-regime Ornstein-Uhlenbeck process (OU1; Hansen, 1997), in which stabilizing selection of strength *α* pulls traits toward an optimum *θ*; (3) bounded BM (Boucher and Démery, 2016), a non-adaptive model with reflective upper and lower bounds; and (4-5) BM with rates that accelerate (AC) or decelerate (DC) exponentially through time at a rate governed by *a* (Blomberg et al., 2003; Harmon et al., 2010b). Parameter distributions are given in Table 1. For each simulated tree–trait dataset, a curated set of twenty summary statistics that describe tree structure, tip-trait distributions, and phylogenetic signal were calculated for use in the neural network (see Tables 2 and 3 for a list of statistics and their definitions).

**Table 1:** Parameter distributions used to simulate trait data. Each parameter was drawn from a uniform distribution, (min, max), with the bounds shown. Parameters are the evolutionary rate (*σ*^2^), selection strength (*α*), selection optimum (*θ*), characteristic time (*τ*), and rate-change parameter (*a*). Dashes indicate parameters not used by a given model. **Alt text** Table 1: Each row represents a candidate evolutionary model considered in this study. Columns represent model parameters and, where relevant, their range of simulated values are defined.

|  | σ<sup>2</sup> | α | θ | τ | *a* |
|---|---|---|---|---|---|
| **BM** | [0, 5] | - | - | - | - |
| **Bounded BM** | [0, 5] | - | - | [0.1, 0.8] | - |
| **OU** | [0, 5] | [0, 5] | [0, 5] | - | - |
| **AC** | [0, 5] | - | - | - | [0, 3] |
| **DC** | [0, 5] | - | - | - | [-3, 0] |

**Table 2:**
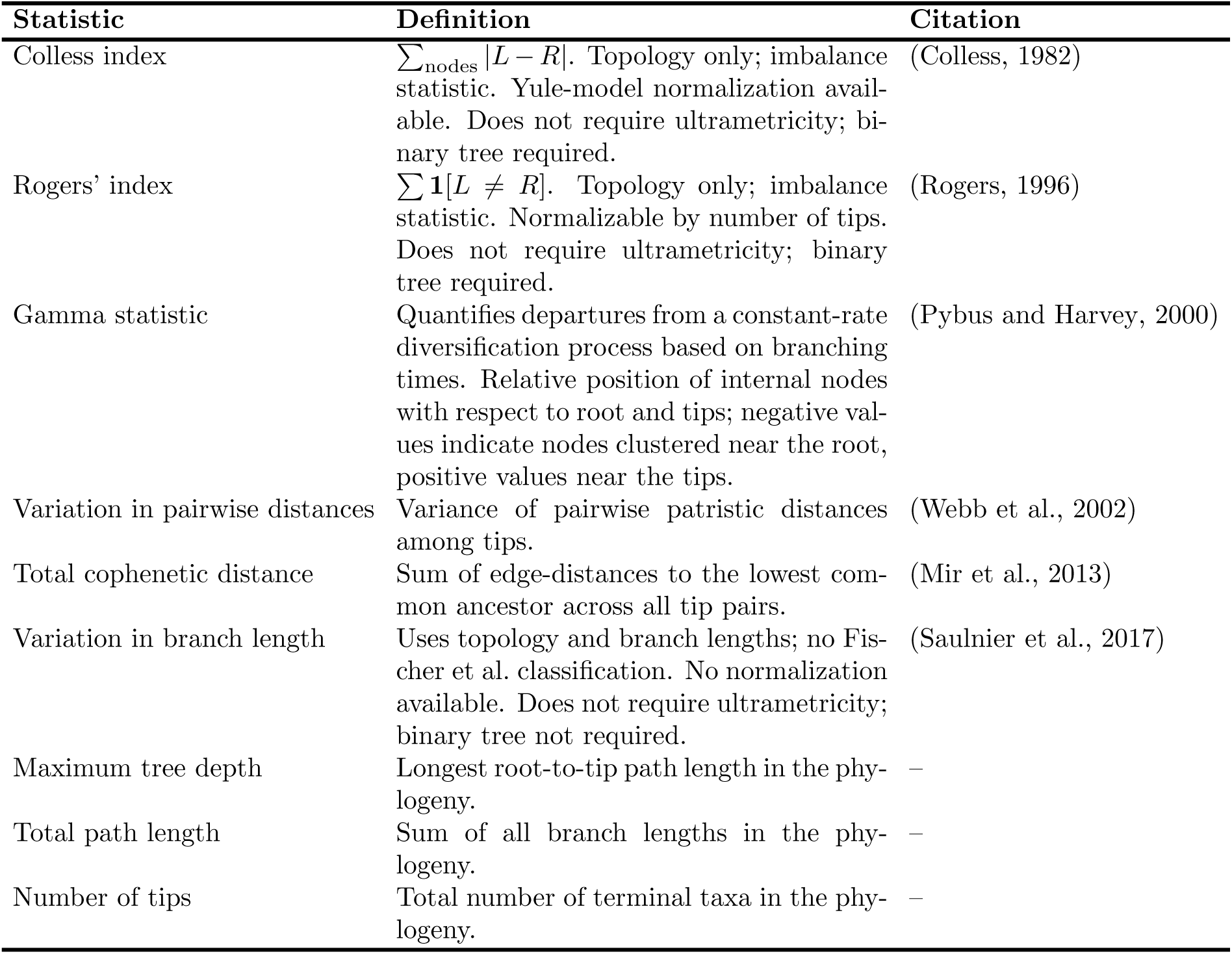
Tree-based summary statistics used as predictor variables in the MLP classification pipeline. **Alt text** Table 2: The nine tree-based summary statistics used in MLP classification. Name, definition, and relevant citations are provided.

| **Statistic** | **Definition** | **Citation** |
|---|---|---|
| Colless index | ∑<sub>nodes</sub> \|*L* − *R*\|. Topology only; imbalance statistic. Yule-model normalization avail-able. Does not require ultrametricity; bi-nary tree required. | (Colless, 1982) |
| Rogers' index | ∑ **1**[*L* ≠ *R*]. Topology only; imbalance statistic. Normalizable by number of tips. Does not require ultrametricity; binary tree required. | (Rogers, 1996) |
| Gamma statistic | Quantifies departures from a constant-rate diversification process based on branching times. Relative position of internal nodes with respect to root and tips; negative val-ues indicate nodes clustered near the root, positive values near the tips. | (Pybus and Harvey, 2000) |
| Variation in pairwise distances | Variance of pairwise patristic distances among tips. | (Webb et al., 2002) |
| Total cophenetic distance | Sum of edge-distances to the lowest com-mon ancestor across all tip pairs. | (Mir et al., 2013) |
| Variation in branch length | Uses topology and branch lengths; no Fis-cher et al. classification. No normalization available. Does not require ultrametricity; binary tree not required. | (Saulnier et al., 2017) |
| Maximum tree depth | Longest root-to-tip path length in the phy-logeny. | – |
| Total path length | Sum of all branch lengths in the phy-logeny. | – |
| Number of tips | Total number of terminal taxa in the phy-logeny. | – |

**Table 3:**
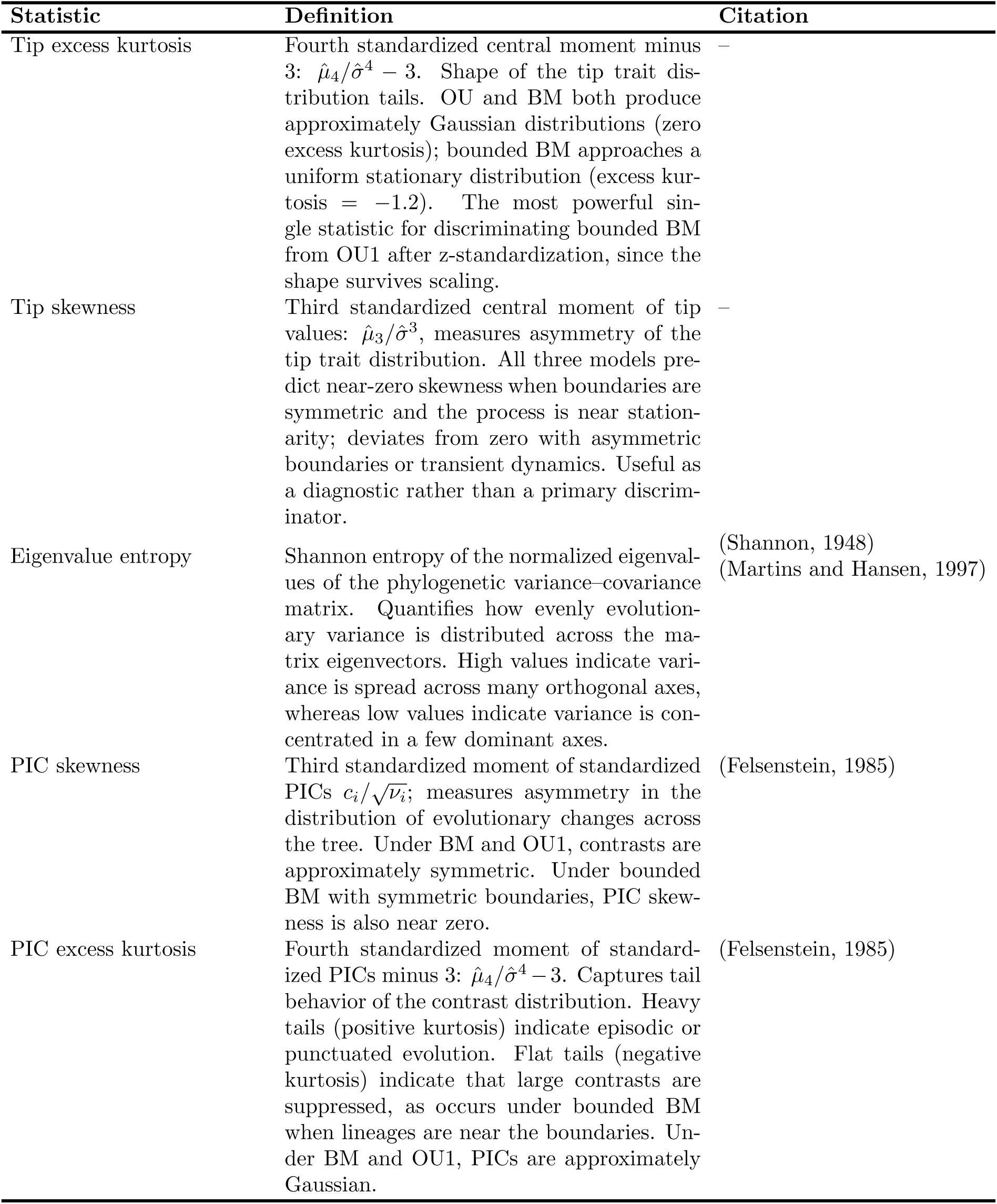
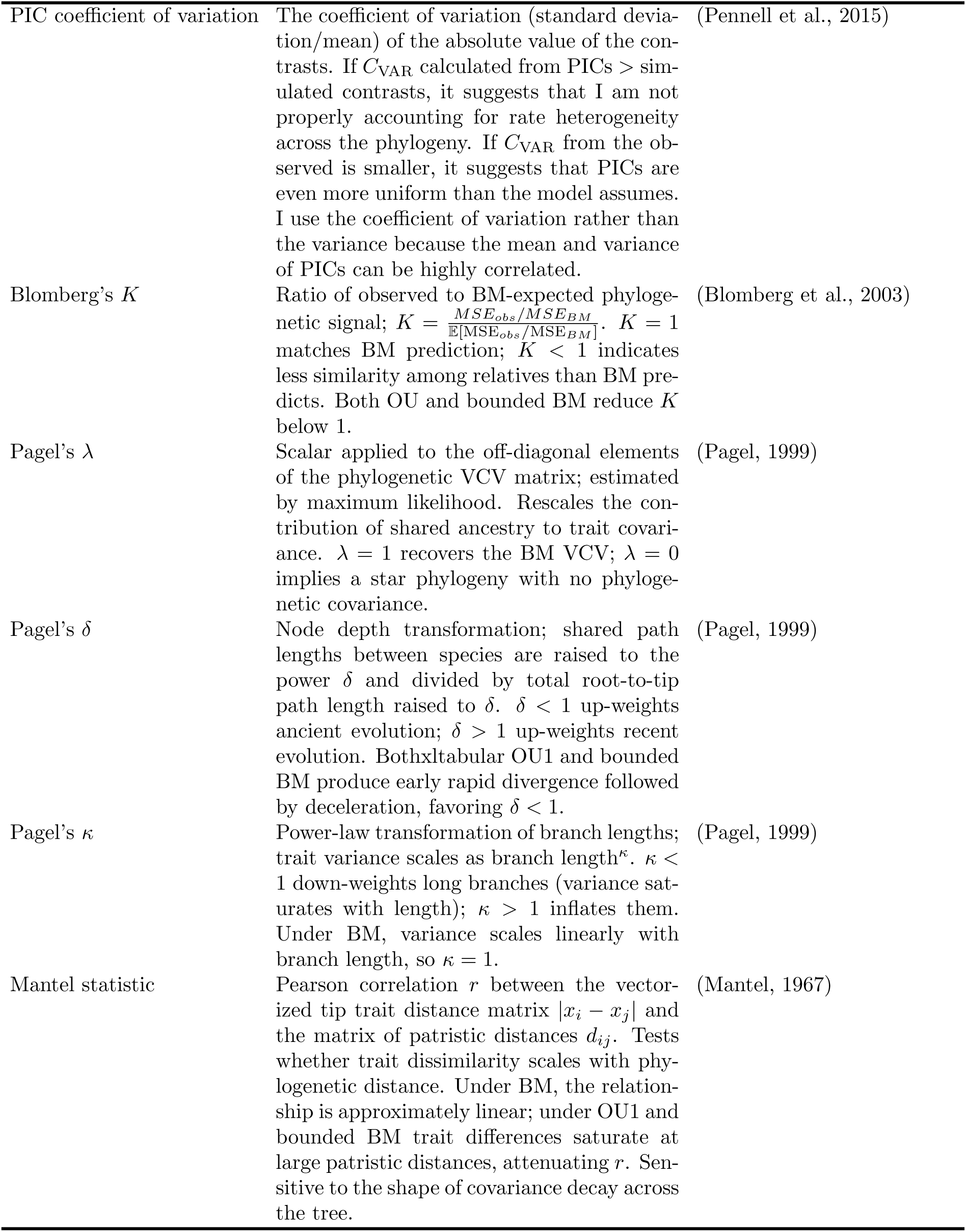
Trait-based summary statistics used as predictor variables in the MLP classification pipeline. **Alt text** Table 3: The eleven trait-based summary statistics used in MLP classification. Name, definition, and relevant citations are provided.

| Statistic | Definition | Citation |
|---|---|---|
| Tip excess kurtosis | Fourth standardized central moment minus 3: μ̂<sub>4</sub>/σ̂<sup>4</sup> − 3. Shape of the tip trait dis-tribution tails. OU and BM both produce approximately Gaussian distributions (zero excess kurtosis); bounded BM approaches a uniform stationary distribution (excess kur-tosis = −1.2). The most powerful sin-gle statistic for discriminating bounded BM from OU1 after z-standardization, since the shape survives scaling. | – |
| Tip skewness | Third standardized central moment of tip values: μ̂<sub>3</sub>/σ̂<sup>3</sup>, measures asymmetry of the tip trait distribution. All three models pre-dict near-zero skewness when boundaries are symmetric and the process is near station-arity; deviates from zero with asymmetric boundaries or transient dynamics. Useful as a diagnostic rather than a primary discrim-inator. | – |
| Eigenvalue entropy | Shannon entropy of the normalized eigenval-ues of the phylogenetic variance–covariance matrix. Quantifies how evenly evolution-ary variance is distributed across the ma-trix eigenvectors. High values indicate vari-ance is spread across many orthogonal axes, whereas low values indicate variance is con-centrated in a few dominant axes. | (Shannon, 1948)<br>(Martins and Hansen, 1997) |
| PIC skewness | Third standardized moment of standardized PICs $c_i/\sqrt{\nu_i}$; measures asymmetry in the distribution of evolutionary changes across the tree. Under BM and OU1, contrasts are approximately symmetric. Under bounded BM with symmetric boundaries, PIC skew-ness is also near zero. | (Felsenstein, 1985) |
| PIC excess kurtosis | Fourth standardized moment of standard-ized PICs minus 3: μ̂<sub>4</sub>/σ̂<sup>4</sup> − 3. Captures tail behavior of the contrast distribution. Heavy tails (positive kurtosis) indicate episodic or punctuated evolution. Flat tails (negative kurtosis) indicate that large contrasts are suppressed, as occurs under bounded BM when lineages are near the boundaries. Un-der BM and OU1, PICs are approximately Gaussian. | (Felsenstein, 1985) |
| PIC coefficient of variation | The coefficient of variation (standard devia-tion/mean) of the absolute value of the con-trasts. If *C*<sub>VAR</sub> calculated from PICs > sim-ulated contrasts, it suggests that I am not properly accounting for rate heterogeneity across the phylogeny. If *C*<sub>VAR</sub> from the ob-served is smaller, it suggests that PICs are even more uniform than the model assumes. I use the coefficient of variation rather than the variance because the mean and variance of PICs can be highly correlated. | (Pennell et al., 2015) |
| Blomberg's *K* | Ratio of observed to BM-expected phyloge-netic signal; $K = \frac{MSE_{obs}/MSE_{BM}}{\mathbb{E}[\mathrm{MSE}_{obs}/\mathrm{MSE}_{BM}]}$. *K* = 1 matches BM prediction; *K* < 1 indicates less similarity among relatives than BM pre-dicts. Both OU and bounded BM reduce *K* below 1. | (Blomberg et al., 2003) |
| Pagel's λ | Scalar applied to the off-diagonal elements of the phylogenetic VCV matrix; estimated by maximum likelihood. Rescales the con-tribution of shared ancestry to trait covari-ance. λ = 1 recovers the BM VCV; λ = 0 implies a star phylogeny with no phyloge-netic covariance. | (Pagel, 1999) |
| Pagel's δ | Node depth transformation; shared path lengths between species are raised to the power δ and divided by total root-to-tip path length raised to δ. δ < 1 up-weights ancient evolution; δ > 1 up-weights recent evolution. Bothxltabular OU1 and bounded BM produce early rapid divergence followed by deceleration, favoring δ < 1. | (Pagel, 1999) |
| Pagel's κ | Power-law transformation of branch lengths; trait variance scales as branch length<sup>κ</sup>. κ < 1 down-weights long branches (variance sat-urates with length); κ > 1 inflates them. Under BM, variance scales linearly with branch length, so κ = 1. | (Pagel, 1999) |
| Mantel statistic | Pearson correlation *r* between the vector-ized tip trait distance matrix \|*x*<sub>*i*</sub> − *x*<sub>*j*</sub>\| and the matrix of patristic distances *d*<sub>*ij*</sub>. Tests whether trait dissimilarity scales with phy-logenetic distance. Under BM, the relation-ship is approximately linear; under OU1 and bounded BM trait differences saturate at large patristic distances, attenuating *r*. Sen-sitive to the shape of covariance decay across the tree. | (Mantel, 1967) |

### Network definition

Given a vector **x** of predictor variables, a feed-forward neural network (FFNN) approximates a function *y*^ = *f* (**x**) that maps a predictor vector **x** to a prediction *y*^, without *a priori* assumptions about functional form (Goodfellow et al., 2016). The vector passes through an input layer, one or more hidden layers, and an output layer. Each hidden neuron computes a weighted sum of the previous layer’s outputs, adds a bias, and applies a nonlinear activation function. During training, weights and biases are iteratively adjusted to minimize a loss function using a backpropagation procedure (Rumelhart et al., 1986; Goodfellow et al., 2016) until a stopping criterion is met. I developed a feed-forward multilayer perceptron (MLP) neural network to classify comparative datasets to their generating evolutionary model: BM, OU1, bounded BM, AC, or DC (Fig. 1). The network input is the vector of summary statistics, which is transformed by the rectified linear unit (ReLU) activation function as it passes through two fully connected hidden layers. A dropout layer is inserted between the hidden layers to reduce overfitting by limiting neuron co-adaptation (Srivastava et al., 2014). The output layer produces a logit score per candidate model that is converted into class probabilities using the softmax function. The evolutionary model assigned to a dataset is the class with the highest predicted probability.

**Figure 1:**
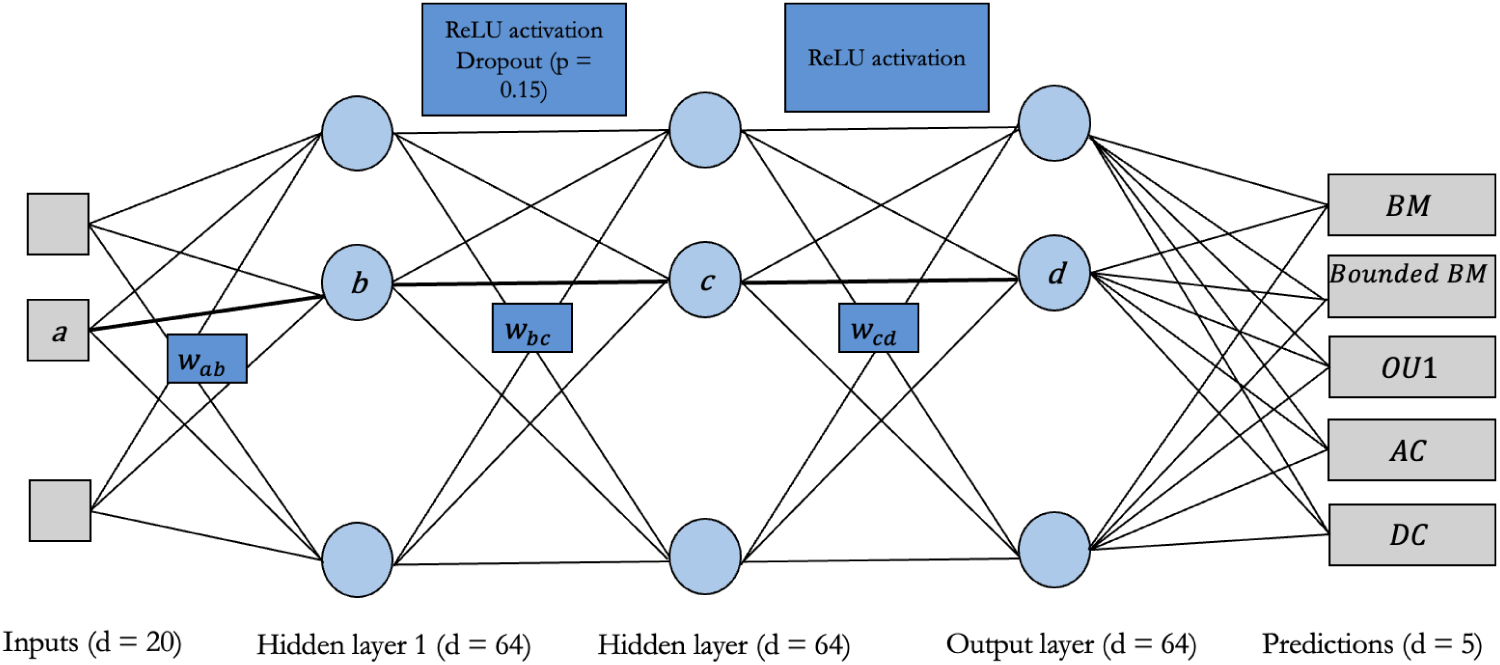
Schematic diagram of the feed-forward multilayer perceptron (MLP) architecture used for model classification. Twenty input summary statistics are passed through two hidden layers for feature extraction. Both hidden layers contains 64 neurons. Each neuron computes a weighted sum of its inputs, adds a bias term, and passes its output through the ReLU activation function. An example neuron-to-neuron pathway is illustrated by nodes *a–b–c–d*, with weights (*w_ij_*) denoting the connection strength between neuron *i* in the previous layer and neuron *j* in the subsequent layer. The network outputs a five-dimensional vector representing the predicted probabilities of the five evolutionary model classes. A dropout rate of 0.15 is applied between the first and second hidden layers for regularization. Adapted from (LeCun et al., 2015), Fig. 1C. **Alt text** Fig.1: Schematic diagram illustrating the key components of the MLP framework. Input and output variables are shown as gray squares. Hidden layers are represent by vertical arrangements of blue circles. Parameter weights of an example input-to-output path-shown in bold - are indicated by the following annotations: *W_AB_, W_BC_, W_CD_*.

### MLP training and validation

The simulated dataset was partitioned 80-10-10 into training, validation, and test subsets (8 × 10^4^, 10^4^, and 10^4^ observations). The training set was used to optimize network parameters, the validation set to monitor performance during training, and the test set was withheld until final evaluation of out-of-sample performance. The MLP was trained by minimizing cross-entropy loss, the standard objective for multi-class classification (Goodfellow et al., 2016), using the Adam optimizer (Kingma and Ba, 2014) on mini-batches of 64 observations (Ruder, 2016). Training ran for a maximum of 500 epochs, with early stopping if validation accuracy did not improve for 50 consecutive epochs. I used K-fold cross validation to optimize the model architecture and training hyperparameters. Full details of this latter step are provided in the Extended Methods section of the Supplementary Material.

### Network evaluation

#### MLP predictive accuracy

I evaluated the MLP’s out-of-sample predictive performance using the test dataset of 10^4^ observations distributed across model classes (*n*_BM_ = 1903, *n*_Bounded_ = 1987, *n*_OU1_ = 2051, *n*_AC_ = 2059, *n*_DC_ = 2000). For direct comparison, MaxL model fitting was performed on the same test datasets using the fitContinuous() function in the geiger package (Harmon et al., 2009) for BM, OU1, AC, and DC models and the bounded bm() function in the phytools package (Revell, 2024) for the bounded BM model. The AC and DC models were treated as separate models by setting bounds on the search space for the *a* parameter of the Early Burst (EB; Blomberg et al., 2003; Harmon et al., 2010b) model such that the rate change parameters were constrained to be positive for AC and negative for DC. I used small sample-corrected AIC weights (Wagenmakers and Farrell, 2004), hereafter AICc weights, as measures of relative support among candidate models and the model receiving the highest weight was selected as best. I constructed confusion matrices to visualize the distribution of prediction accuracy and misclassifications across model classes for MLP-and MaxL-based predictions.

#### Comparing model discriminability under ML and MaxL

Likelihood-based evolutionary model selection is negatively impacted by small phylogenies and the magnitude of generative parameters (Revell et al., 2008; Münkemüller et al., 2012; Boettiger et al., 2012; Ho and Ané, 2014; Cooper et al., 2016b). To assess whether the MLP exhibits similar sensitivity, I evaluated classification accuracy as a function of tree size and generating parameter values for both MaxL and MLP classifications using Bayesian beta regression with a logit link, fitted via the brms package (Bürkner, 2017) (see the Extended Methods section of the Supplementary Material for full details). Thin-plate regression splines were used to accommodate nonlinearity in the marginal effects of each predictor (Wood, 2003). For models with two or more generating parameters (bounded BM, OU1, AC, DC), tensor product smooths were constructed for each parameter-tree size pairing to examine how the influence of each generating parameter on the outcome varied as a function of tree size (Wood, 2003).

#### Evaluating variable importance in MLP classification using BART

It can be difficult to understand which aspects of a given dataset contribute most to successful prediction in neural network approaches. I used Bayesian Additive Regression Trees (BART) to better understand the contribution of summary statistics to the identification of the five candidate models. BART is a nonparametric ensemble ML method that models complex, nonlinear relationships between predictor and outcome variables through a sum of regression trees (Chipman et al., 2010) that recursively partition the predictor space through binary splits and assign associated parameter values to terminal nodes. Using functions in the bartMachine package (Kapelner and Bleich, 2016), I trained BART models for each evolutionary model class, considering the summary statistics as predictor variables and the logit-transformed MLP probability assigned to the correct model class as the response variable. Each dataset was randomly divided into training (80%) and testing (20%) subsets, with model hyperparameters chosen using crossvalidation across ensemble sizes (*m* = 10, 20, 30), and regularization strength parameters (*k* = 1, 2, 3) using the bartMachineCV function. Predictive performance was evaluated using root mean squared error (RMSE) and mean absolute error (MAE) calculated on held-out test data after transforming predictions back to probability space and final models selected on the combination of predictive performance and model stability. Variable importance was quantified using the investigate var importance() function, which measures the proportion of times each variable is chosen for a splitting rule across the ensemble. A more complete description of model construction and evaluation is provided in the Extended Methods section of the Supplementary Methods.

### Empirical case study: Detecting signals of early burst dynamics in comparative data

G.G. Simpson (1953) famously credited adaptive radiations – explosive morphological diversification that slows through time – with explaining the origin of most of life’s diversity. Harmon et al. (2010b) tested how common such early bursts (EBs) are by asking how often their signal could be detected in comparative data, including body size and shape, from 49 clades (Harmon et al., 2010b, Table 1). They found EB (=DC) support was strikingly rare, BM was the model most frequently preferred in clades with more than 10 tips, and a Single Stationary Peak (SSP=OU1) was the best single model based on pooled likelihoods (Harmon et al., 2010b), leading them to conclude that EBs are rare in comparative data. I hypothesized that this lack of support could reflect a failure to detect EBs in comparative data using Likelihood methods, rather than a scarcity of early burst events. If this hypothesis is correct, then MLP classifications should assign more support to the EB model than Harmon et al. (2010b) did, largely at the expense of BM. I further hypothesized that, in some cases, preference for SSP in Harmon et al. (2010b) reflects the absence of alternative candidate models that produce similarly low phylogenetic signal, notably bounded BM. If true, then the MLP should also assign less support to SSP and redistribute some to bounded BM.

To test these hypotheses, I obtained data from the online supplementary material of Harmon et al. (2010a). Because the MLP was trained on trees of 20 to 1000 tips, I retained only datasets with at least 20 sampled species to avoid extrapolation. The final set consisted of 29 datasets (18 body size, 11 body shape) spanning taxonomic resolutions from genus to suborder level. To directly compare MLPand MaxL-based classifications, I refit BM, SSP, and EB to the datasets as well as a bounded BM candidate model. I avoided the non-identifiability of AC and OU1 (Uyeda et al., 2015) by retraining the MLP omitting AC and with OU1 and DC relabeled as SSP and DC to match Harmon et al. (2010b). For each tree-trait dataset, I rescaled the tree to unit depth and calculated summary statistics, which I standardized using the means and standard deviations of the simulated training set. This places empirical data on the scale on which the network was trained, and avoids recalculating scaling parameters for each dataset. The standardized statistics were passed to the trained MLP, which returned a probability for each model in every dataset. Finally, I compared the MLP class probabilities with the AIC weights published by Harmon et al. (2010b) and with AIC weights from refitting the expanded candidate set.

## Results

### How useful is the MLP for classifying comparative data?

#### Is the classifier accurate?

MaxL model selection achieved an overall predictive accuracy of 54.8% across the 10,000 simulated observations, where accuracy is defined as the highest AICc weight belonging to the generating process, regardless of magnitude. Performance varied among model classes, with high accuracy for bounded BM (100%) and OU1 (82%) models, but poor accuracy for BM (34%) and DC (26%) (Fig. 2). I found a complete lack of true AC model discrimination (0%), with support instead assigned to OU1 (55%), bounded BM (35%), and BM (10%). The MLP achieved an overall accuracy of 74.3% on the test dataset and performed more consistently than MaxL across model classes (Fig. 2). All five classes were correctly identified for the majority of simulations (all diagonals in Fig. 2B *>* 0.5), with per-class accuracy ranging from 64% for OU1 and AC to 100% for bounded BM models. Notably, BM is correctly predicted in over 75% of cases under MLP, compared with only 34% under MaxL classification. This represents a genuine preference for the BM model instead of a mere failure to reject it in favor of alternative models. When misclassification did occur, it was typically in the form of assigning true AC and DC cases to BM. Conflation of AC with OU1 occurred in only 7% of cases, representing a large reduction relative to the 55% misassignment observed under MaxL model selection, though the reverse case of OU1 misidentified as AC rose from 0% under MaxL model selection to 26% under MLP (but see below). The four-class MLP trained for empirical application achieved a higher overall test accuracy relative to the original five-class MLP (85.4%, Fig. S1). Class-specific accuracy ranged from 69% for EB to 100% for bounded BM. The most common errors were misclassifying EB as BM in 30% of true EB cases and SSP as BM in 14% of true SSP cases.

**Figure 2:**
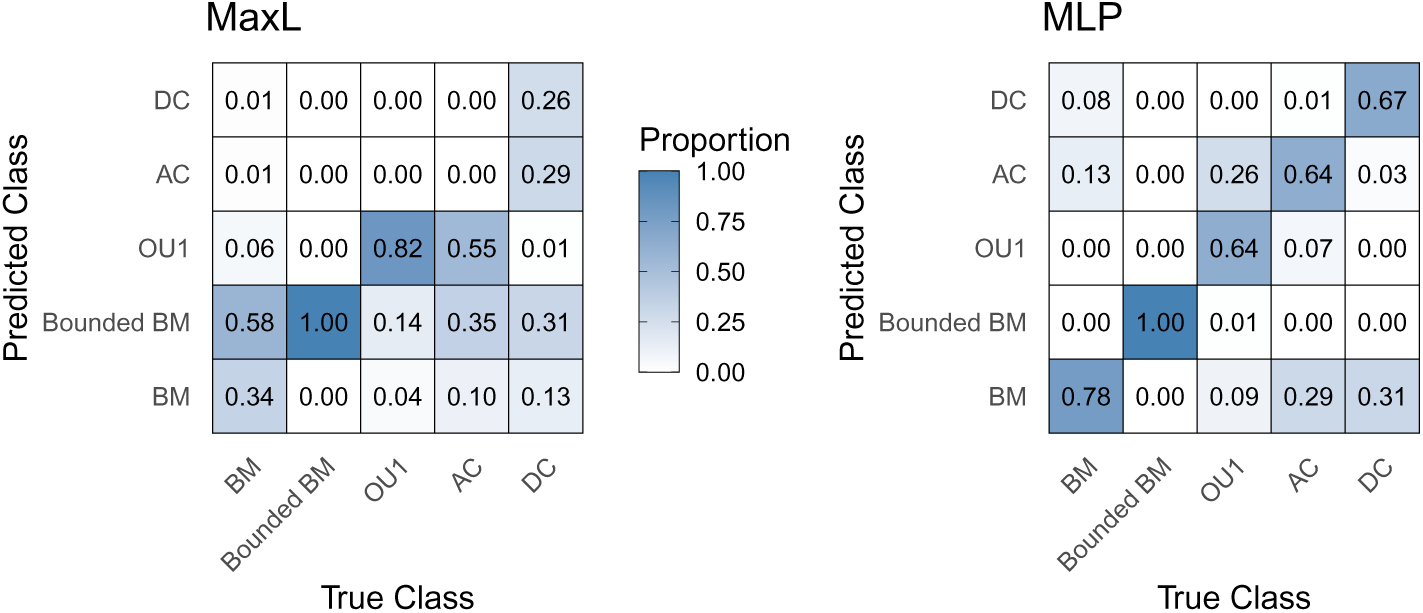
Confusion matrices to visualize the distribution of classifications for traditional MaxL model fitting (left) and the MLP pipeline (right). Rows represent the true model class and columns show the predicted class such that for each column, proportions total to one. **Alt text** Fig.2: Distribution of evolutionary model classification errors for both the MaxL (left) and MLP (right) classifier frameworks. Rows denote the true model class, columns denote the predicted model class. Each cell contains the proportion of times, for each true model class, that model is predicted across the test data frame. White tiles received no support, dark blue tiles received (near-)full support.

#### Is the classifier biased?

Under MaxL, baseline accuracy (the posterior predicted probability of correct classification at mean tree size and parameter values) varied considerably among models (Fig. 3; Table S2). It was near-perfect for bounded BM, rendering parameter and tree size effects uninformative, but poor for BM (22%), AC (3.5%), and DC (36%), and higher for OU1 (77%). OU1 performance rose nonlinearly with *α* to a plateau at intermediate values, sharpest for the 90th-percentile tree size (Fig. 3A). AICc weights for a true AC model remained low across all values of *a*, declining slightly as *a* increased, with curves indistinguishable among tree sizes (Fig. 3B). The MLP also achieved near-perfect baseline accuracy for bounded BM (*>* 99.7%) and variable accuracy for the other models (BM: 47%, OU1: 73%, AC: 45%, DC: 69%; Table S2). In contrast with MaxL, tree size curves for the MLP were similar, with the 50th-and 90th-percentile curves largely indistinguishable, indicating a weak effect of tree size. OU1 performance again rose toward a plateau with *α* (Fig. 3D). For AC, the MLP showed a stronger relationship with *a* than MaxL (Fig. 3E): for medium and large trees, the probability assigned to AC peaked at intermediate values (*a* ≈ 2.1, 95% CI: 1.94-2.21; and *a* ≈ 2.0, 95% CI: 1.81-2.12, respectively) and declined thereafter. For DC, performance under both classifiers increased as *a* became more negative, peaking at the lower boundary (*a* = −3) and reaching its lowest at the upper boundary (*a* = 0) (Fig. 3C,F).

**Figure 3:**
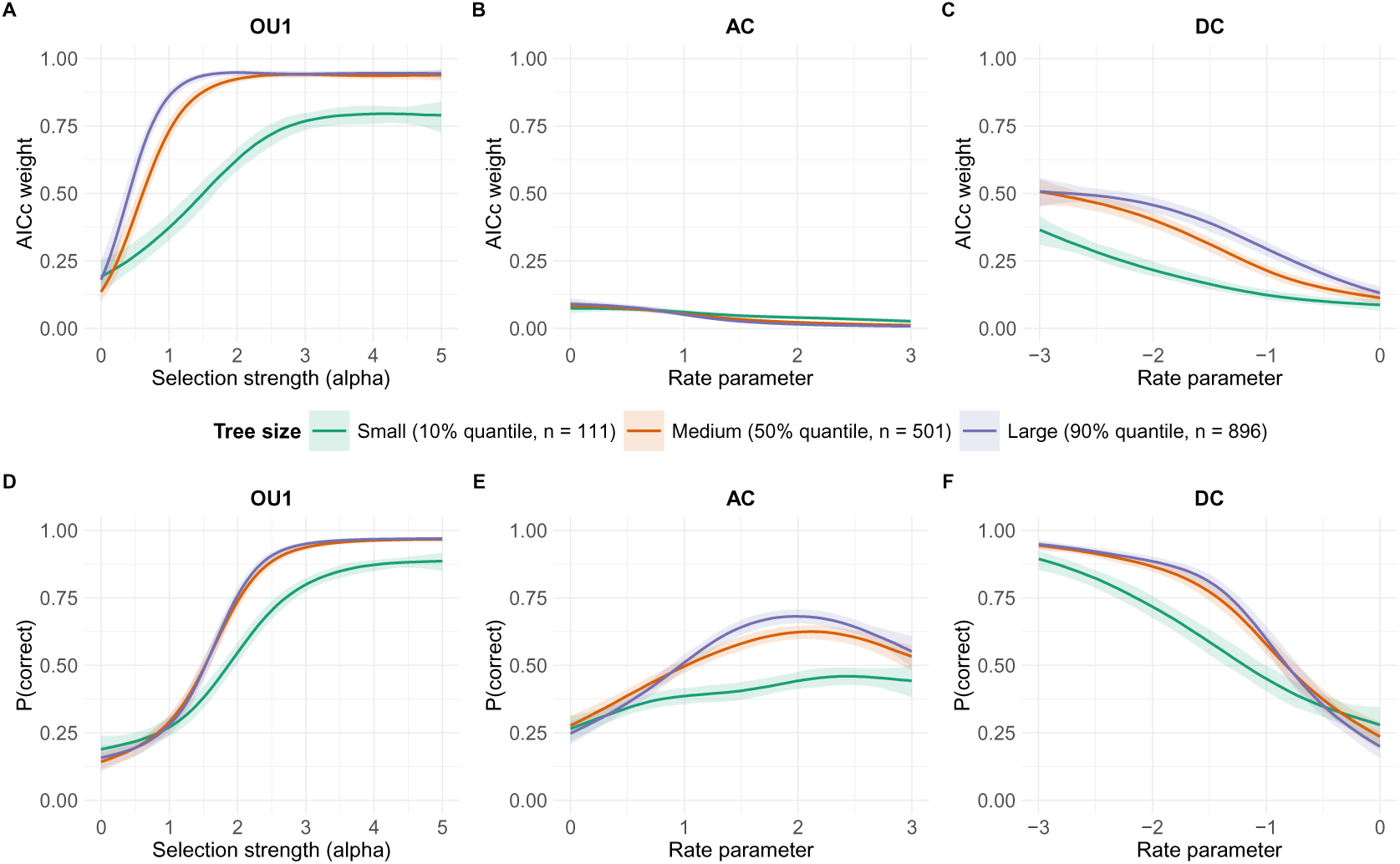
Posterior predictions of the AICc weight (A-C) and probability (D-F) assigned to the true generating model as a function of generative parameters, displayed across the 10th, 50th, and 90th percentiles of the tree size distribution. A-C show predictions from MaxL estimation; D-F show predictions from the MLP classifier. Predictions are generated using Bayesian beta regression with a logit link. Shaded intervals represent 95% confidence bands. **Alt text** Fig.3: Six-panel figure showing posterior predictions of the support for the true evolutionary model class across the test data frame. Panels A to C: AICc weight assigned to the true generating model by MaxL classification. Panels D to F: probability assigned to the true generating model by MLPbased classification. For each panel, the three curves represent different tree size percentiles: 10th (teal), 50th (orange), and 90th (purple). Generative parameter values (x-axis) are plotted against the model support metric (y-axis). Shaded intervals represent 95% confidence bands.

### What drives the predictive performance of the MLP?

I used BART models to identify variables associated with successful classification to the generating model. Predictor importance, measured as inclusion proportion, shows that some summary statistics are important for all models, while others aid in the discrimination of specific classes (Fig. 5, Table S4). Blomberg’s *K* is recovered in the top 2 most important predictors for BM, OU1, AC, and DC, making it the single most useful predictor, though it does not alone discriminate any one model. Pagel’s *κ* and *δ* also feature prominently, though their order of importance varies among models. Trait-based summary statistics are consistently included more frequently in the “above random” set than tree-based statistics. BM is discriminated by a strong effect of *δ*, followed by *K* and *κ*. OU1 detection also relies on *K* and *κ* but *δ* is reduced to near the bottom of the list. AC and DC both rely on *δ* and *K*, though the order of importance flips and their secondary predictors are different; AC relies on eigenvalue entropy and *κ* while DC on the Mantel statistic for trait and patristic distances and the PIC excess kurtosis. Bounded BM has the most unusual profile. The variable inclusion proportion distribution is nearly flat, and it is the only process for which tip excess kurtosis ranks highly, while *δ* and *κ*, which are so important to the other four models, ranks near the bottom.

### OU1-AC equivalence

My results (Figure 2) imply that AC-OU1 discrimination is possible using the MLP. However, on ultrametric trees, AC and OU1 models are non-identifiable because each OU1 parameter value (*α*) has a corresponding AC parameter (*a*) that yields an identical likelihood, specifically *a* = 2*α* (Uyeda et al., 2015). Because MLP can only learn patterns from the data on which it is trained, fair evaluation of AC-OU1 discrimination therefore requires testing in comparable parameter spaces across the simulated models, which is not the case here: the maximum simulated *α* = 5 is equivalent to *a* = 10, which lies beyond the simulated AC range (*a* ≤ 3). Additionally, for true AC, the tip rate, *σ*^2^, must lie within the range of *σ*^2^ simulated under OU1 (0 *< σ*^2^ *<* 5). I therefore performed a second comparison in which I restricted the test data to AC and OU1 simulations in the equivalence space (OU1 *α* ≤ α_max_/2; AC *σ*^2^_0_ ≤ σ^2^_OU1_*e^−a^*). The resulting subset contained 1272 observations, of which 47.7% were OU1 and 52.3% were AC. I then plotted the MLP-derived *P* (AC) and *P* (OU1) against the generating parameter of each model (*a* for AC, 2*α* for OU1), using loess smoothing. Identical curves for each model probability across both generating models would indicate that the two models are indistinguishable in the equivalence region, whereas differences would suggest that the MLP detects genuine differences in trait summary statistics that are not captured by likelihood metrics. The smoothed curves are almost identical (Fig. 4) and E[*P* (AC)] exceeded E[*P* (OU1)] at all parameter values, regardless of generating model, confirming that these models remain non-identifiable under MLP. The high prediction accuracy for AC appears to arise from arbitrary tie-breaking in favor of AC in the shared parameter space, whereas its high accuracy for OU1 arises from training on and testing with OU1 simulations outside this shared space.

**Figure 4:**
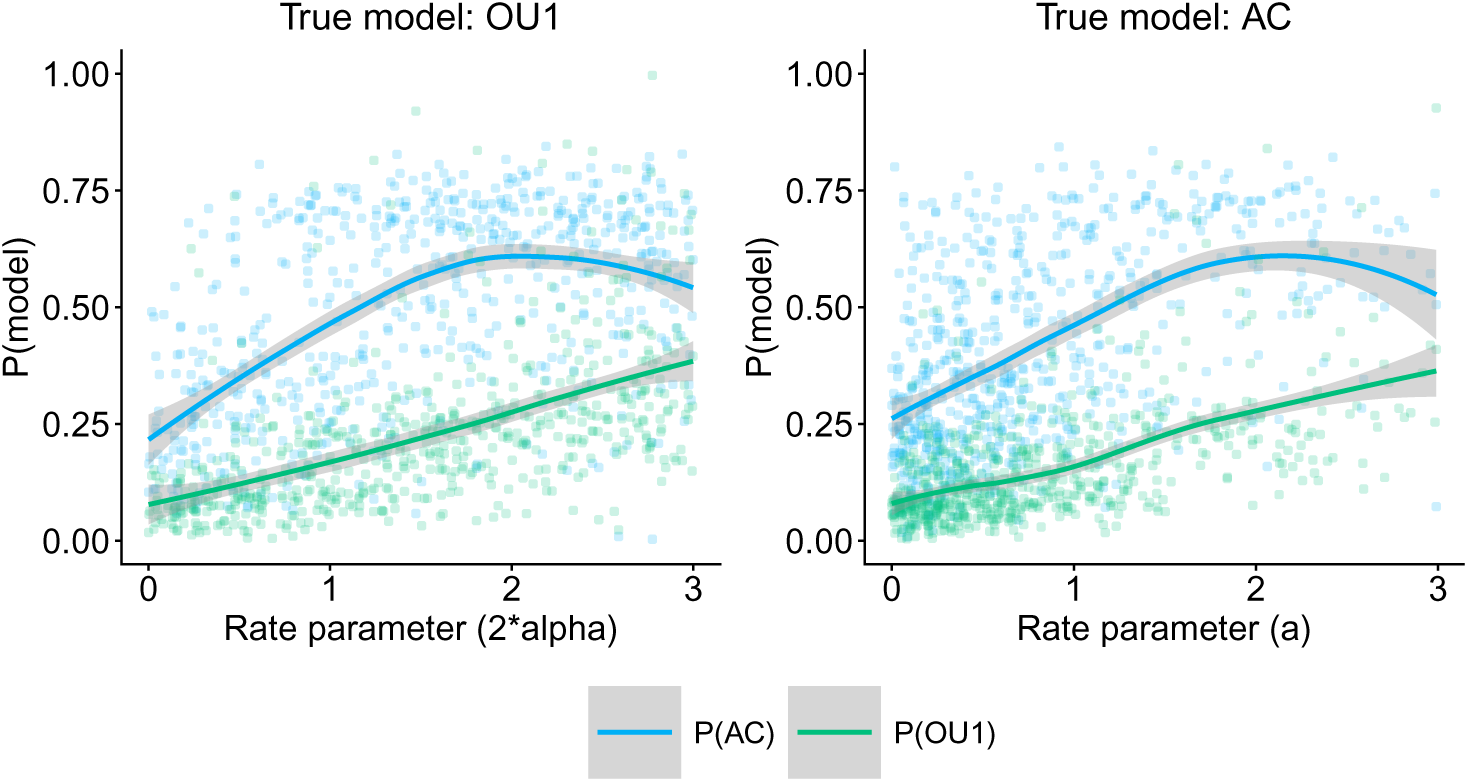
Probabilities assigned by the MLP to AC and OU1 as a function of the rate parameter, within the likelihood-equivalent region of parameter space: 2*α* 3 for OU1; *σ*^2^ 5 exp(*a*) for AC (Uyeda et al., 2015) Panel A displays simulations generated under OU1 (rate parameter: 2*α*) and panel B displays simulations generated under AC (rate parameter: *a*). Individual curves represent loess-smoothed probabilities assigned to AC and to OU1, with shaded bands showing the 95% confidence interval bands. The near-identical curves across the two panels suggests that the MLP fails to discern AC-OU1 models in the same region where likelihood equivalence holds true. **Alt text** Fig. 4: Two panels displaying the probability the MLP assigns to AC and to OU1 against the rate parameter: 2 ∗ *α* for OU1; *a* for AC. The left panel (A) shows data simulated under OU1 and the right panel (B) shows data simulated under AC. Each panel contains two separate smoothed curve for the probability assigned to AC and OU1, with a shaded 95% confidence region. P(AC) exhibits a curved form and peaks at *P* (*AC*) = 0.61*, a* = 2.04 for a true OU1 and *P* (*AC*) = 0.61*, a* = 2.15 for a true AC model, before beginning to decline. P(OU) increases in a consistent but non-linear fashion with the rate parameter across the two models and peaks at *P* (*OU* 1) = 0.38*, alpha* = 1.50 and *P* (*OU* 1) = 0.36*, alpha* = 1.50 for a true OU1 and AC model, respectively. The curves have similar forms across panels, to reflect consistent probability assignments by the MLP, regardless of true generative model.

### Empirical case study: Detecting signals of early burst dynamics in comparative data

Harmon et al. (2010b) assessed the frequency of EBs in comparative data using MaxL and AIC*_c_* comparison of three models: BM, SSP (= OU1), and EB (= DC). I revisited this study using 29 datasets with *N* tips ≥ 20. The original analysis yielded AIC*_c_* weights that favored BM in 19 datasets, SSP in nine, and EB in only one. Repeating model fitting with bounded BM added to the candidate set resulted in AIC*_c_*weights that favored bounded BM in 20 of the 29 datasets (Fig. 6, Tables 4 and 5), with full weight assigned to bounded BM for body size evolution in the Old World primates and Phrynosomatinae. Among the remaining nine datasets, SSP was favored in seven, and BM and EB in one each. The single EB dataset was the same one that received dominant support in the original analysis (*Varanus* body size), although its AIC weight was slightly lower (0.533 versus 0.624).

**Figure 5:**
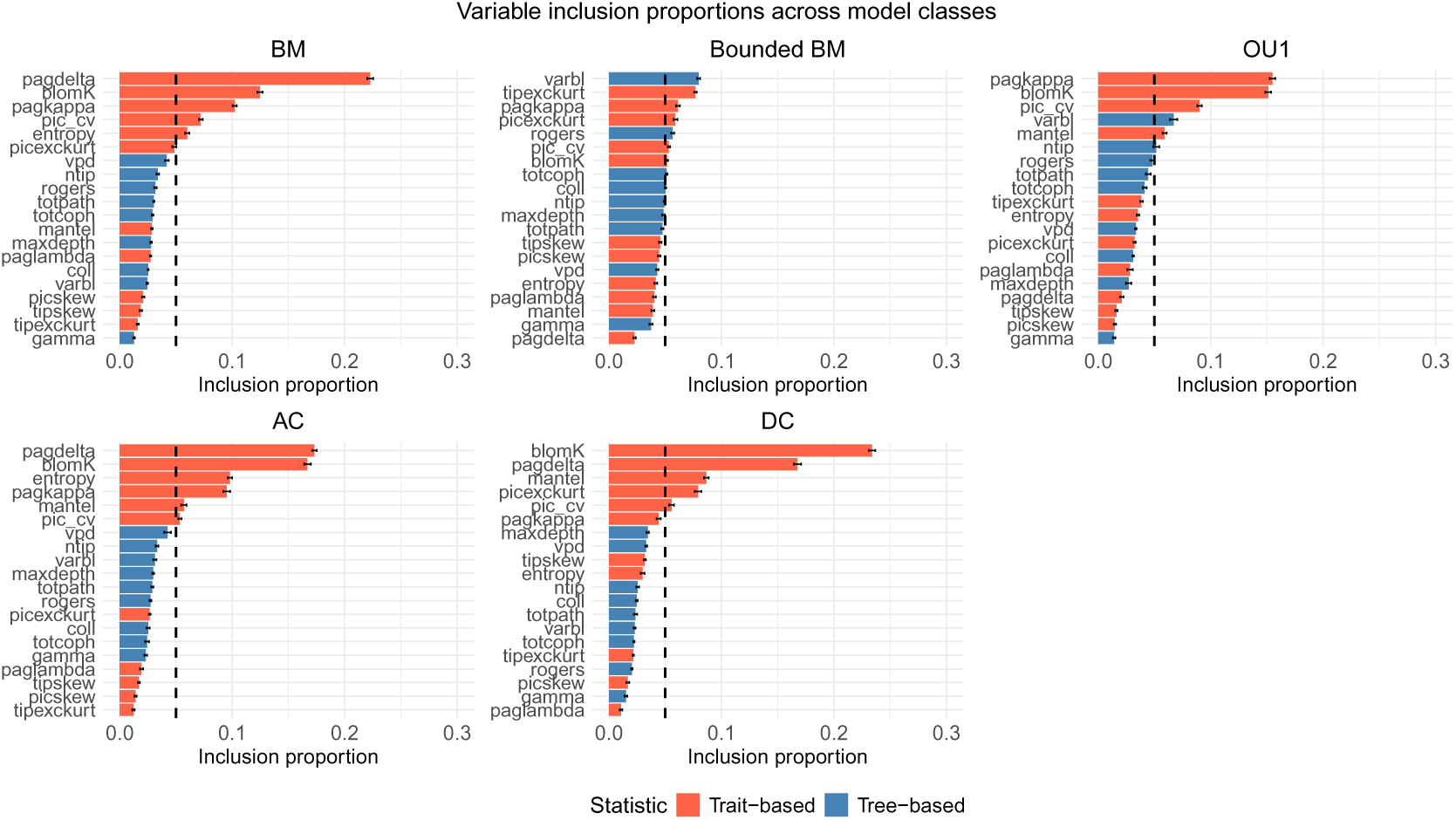
Variable inclusion plots for each BART model generated from 50 replicates. A greater inclusion proportion indicates greater influence on splitting rules within each individual decision tree across the ensemble. Dashed line shows expected included proportions if all variables are independent of the outcome (classification). Black bars represent the mean estimate +*/* the standard deviation across simulation replicates. Colors indicate treeversus trait-based predictors. Variable abbreviations: *pagdelta*, Pagel’s *δ*; *blomK*, Blomberg’s *K*; *pagkappa*, Pagel’s *κ*; *pic cv*, PIC coefficient of variation; *entropy*, eigenvalue entropy; *picexckurt*, PIC excess kurtosis; *vpd*, variation in pairwise distances; *ntip*, number of tips; *rogers*, Rogers’ index; *totpath*, total path length; *totcoph*, total cophenetic distance; *mantel*, Mantel statistic; *maxdepth*, maximum tree depth; *paglambda*, Pagel’s *λ*; *coll*, Colless index; *varbl*, variation in branch length; *picskew*, PIC skewness; *tipskew*, tip skewness; *tipexckurt*, tip excess kurtosis; and *gamma*, *γ* statistic. **Alt text** Fig. 5: Variable inclusion plots for the MLP classification of each evolutionary model class used in training. Values on the x-axis represent the proportion of splitting nodes in which each individual predictor contributed to the splitting rule. Trait-based statistics are shown in red and treebased statistics are shown in blue. The vertical dashed line shows the expected inclusion proportions under the null model of no predictor-outcome relationship. Black bars represent the mean estimate +*/*− the standard deviation across simulation replicates.

**Figure 6:**
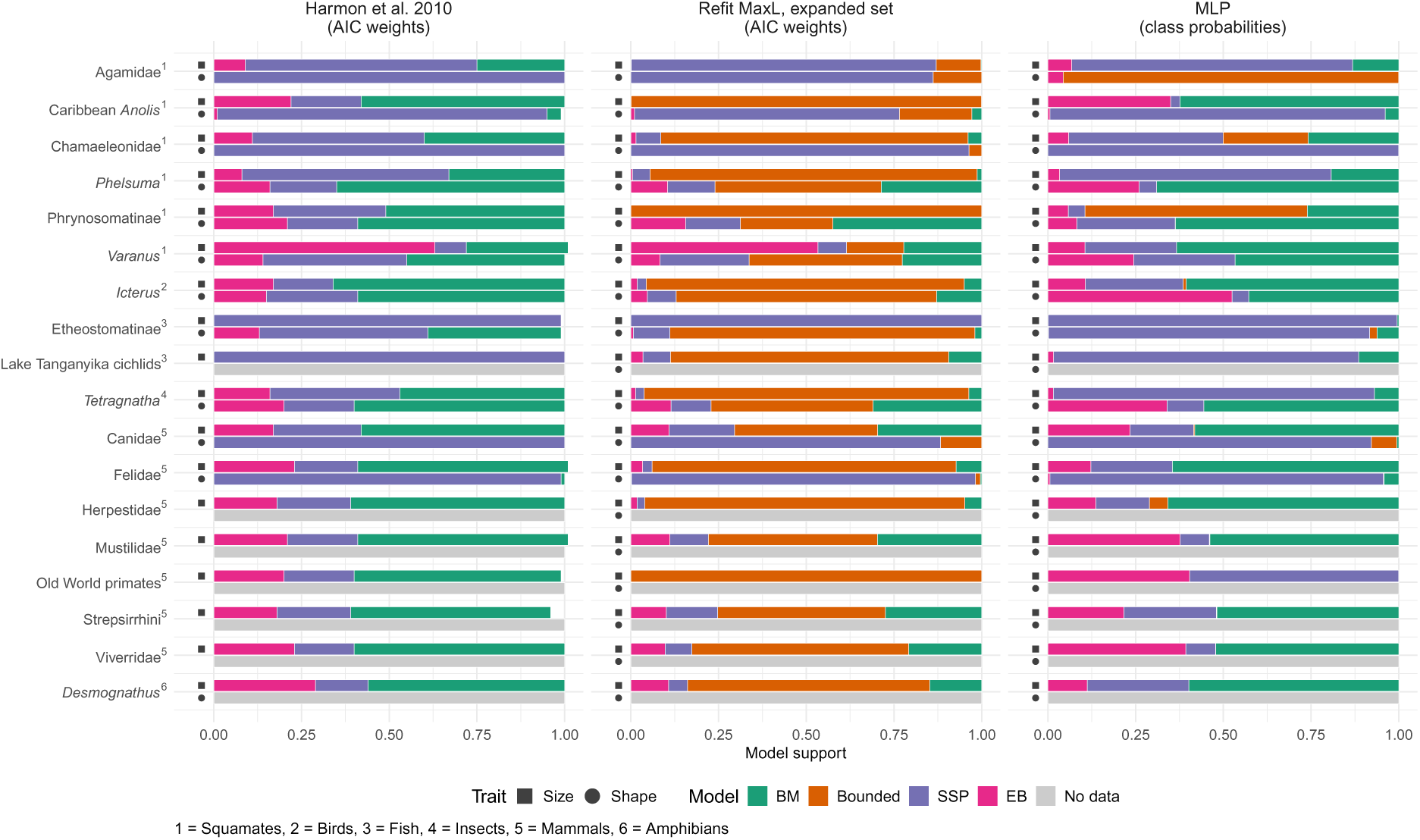
Model support for body size and body shape evolution in the datasets of Harmon et al. (2010b), under maximum likelihood (MaxL) model fitting and MLP classification. Left: AIC weights published by Harmon et al. (2010b) for Brownian motion (BM), Single Stationary Peak (SSP) and Early Burst (EB) models. Right: class probabilities from the four-class MLP, which includes an additional bounded BM model class. Each dataset has two stacked bars whose segments sum to one: body size (upper bar, square) and body shape (lower bar, circle). Grey bars indicate that no body shape data were available. **Alt text** Fig. 6: Two-panel stacked bar chart comparing model support for body size and body shape evolution across 18 clades. The left panel shows AIC weights published by Harmon et al. (2010b) and the right panel shows class probabilities generated by the trained four-class MLP. Each bar is divided into colored segments for BM, bounded BM, SSP and EB. Each clade has an upper bar for body size (square) and a lower bar for body shape (circle). Grey bars mark clades with no body shape data. In the left panel, BM has the largest share in 12 of the 18 body size bars. In the right panel, SSP has the largest share in more datasets and the additional bounded BM model is favored in the body size bar of Phrynosomatinae and the body shape bar of Agamidae.

**Table 4:** Comparison of model fitting analyses using likelihood-based AIC weights (Harmon et al., 2010b), AIC weights from refitting an expanded candidate set that included bounded BM, and MLP classification probabilities, fitted to quantify **body size** evolution. AIC weights from Harmon et al. (2010b) compared three models: BM, SSP, and EB. Refit AIC weights and MLP probabilities covered four models: BM, SSP, EB, and bounded BM. *n* is the number of species (tips) in each dataset. For each set of analyses, the dominant model is indicated in bold. All data is taken from the supplement material of Harmon et al. (2010b). **Alt text** Table 4: Model support for body size evolution in each dataset, shown as likelihood-based AIC weights from Harmon et al. (2010b), AIC weights from refitting an expanded candidate set including bounded BM, and MLP classification probabilities for BM, SSP, EB and bounded BM. The number of species is given for each dataset, and the dominant model in each set of analyses is in bold.

| Dataset | *n* | AIC weights |  |  | Refit AIC weights |  |  |  | MLP probabilities |  |  |  |
|---|---|---|---|---|---|---|---|---|---|---|---|---|
|  |  | BM | SSP | EB | BM | SSP | EB | Bounded | BM | SSP | EB | Bounded |
| Agamidae | 179 | 0.25 | **0.66** | 0.09 | 2.44 × 10<sup>−3</sup> | **0.869** | 8.92 × 10<sup>−4</sup> | 0.128 | 0.132 | **0.801** | 0.0678 | 2.05 × 10<sup>−5</sup> |
| Canidae | 33 | **0.58** | 0.25 | 0.17 | 0.297 | 0.186 | 0.109 | **0.408** | **0.592** | 0.181 | 0.224 | 3.20 × 10<sup>−3</sup> |
| Caribbean *Anolis* | 73 | **0.58** | 0.2 | 0.22 | 7.59 × 10<sup>−4</sup> | 2.79 × 10<sup>−4</sup> | 3.09 × 10<sup>−4</sup> | **0.999** | **0.624** | 0.0266 | 0.350 | 6.46 × 10<sup>−8</sup> |
| Chamaeleonidae | 29 | 0.4 | **0.49** | 0.11 | 0.0390 | 0.0708 | 0.0144 | **0.876** | 0.258 | **0.440** | 0.0592 | 0.242 |
| Etheostomatinae | 122 | 0 | **1** | 0 | 3.02 × 10<sup>−5</sup> | **1.00** | 1.11 × 10<sup>−5</sup> | 1.49 × 10<sup>−4</sup> | 0.00459 | **0.995** | 2.59 × 10<sup>−4</sup> | 2.87 × 10<sup>−8</sup> |
| *Desmognathus* | 23 | **0.56** | 0.15 | 0.29 | 0.148 | 0.0545 | 0.107 | **0.690** | **0.599** | 0.289 | 0.112 | 3.11 × 10<sup>−5</sup> |
| Felidae | 36 | **0.594** | 0.178 | 0.228 | 0.0732 | 0.0269 | 0.0335 | **0.866** | **0.645** | 0.232 | 0.123 | 1.21 × 10<sup>−4</sup> |
| *Tetragnatha* | 59 | **0.47** | 0.37 | 0.16 | 0.0367 | 0.0245 | 0.0135 | **0.925** | 0.0695 | **0.915** | 0.0154 | 6.20 × 10<sup>−7</sup> |
| Lake Tanganyika cichlids | 76 | 0 | **1** | 0 | 0.0943 | 0.0780 | 0.0347 | **0.793** | 0.114 | **0.869** | 0.0162 | 5.21 × 10<sup>−5</sup> |
| Herpestidae | 31 | **0.61** | 0.21 | 0.18 | 0.0482 | 0.0219 | 0.0177 | **0.912** | **0.642** | 0.204 | 0.111 | 0.0436 |
| *Icterus* | 23 | **0.66** | 0.17 | 0.17 | 0.0500 | 0.0256 | 0.0184 | **0.906** | **0.606** | 0.279 | 0.107 | 8.69 × 10<sup>−3</sup> |
| Mustilidae | 62 | **0.594** | 0.198 | 0.208 | 0.297 | 0.109 | 0.112 | **0.482** | **0.567** | 0.0887 | 0.343 | 1.57 × 10<sup>−3</sup> |
| Old World primates | 89 | **0.596** | 0.202 | 0.202 | 2.57 × 10<sup>−17</sup> | 8.16 × 10<sup>−13</sup> | 9.46 × 10<sup>−18</sup> | **1.00** | 1.61 × 10<sup>−4</sup> | **0.596** | 0.404 | 4.48 × 10<sup>−14</sup> |
| *Phelsuma* | 20 | 0.33 | **0.59** | 0.08 | 0.0128 | 0.0501 | 4.65 × 10<sup>−3</sup> | **0.932** | 0.192 | **0.774** | 0.0334 | 6.96 × 10<sup>−7</sup> |
| Phrynosomatinae | 70 | **0.51** | 0.32 | 0.17 | 5.71 × 10<sup>−5</sup> | 4.49 × 10<sup>−5</sup> | 2.10 × 10<sup>−5</sup> | **1.00** | 0.261 | 0.0472 | 0.0583 | **0.634** |
| Strepsirrhini | 44 | **0.594** | 0.219 | 0.188 | 0.274 | 0.146 | 0.101 | **0.478** | **0.518** | 0.264 | 0.216 | 1.37 × 10<sup>−3</sup> |
| *Varanus* | 38 | 0.287 | 0.0891 | **0.624** | 0.222 | 0.0816 | **0.533** | 0.163 | **0.633** | 0.261 | 0.106 | 3.83 × 10<sup>−7</sup> |
| Viverridae | 30 | **0.6** | 0.17 | 0.23 | 0.208 | 0.0765 | 0.0976 | **0.618** | **0.522** | 0.0850 | 0.393 | 2.97 × 10<sup>−4</sup> |

**Table 5:**
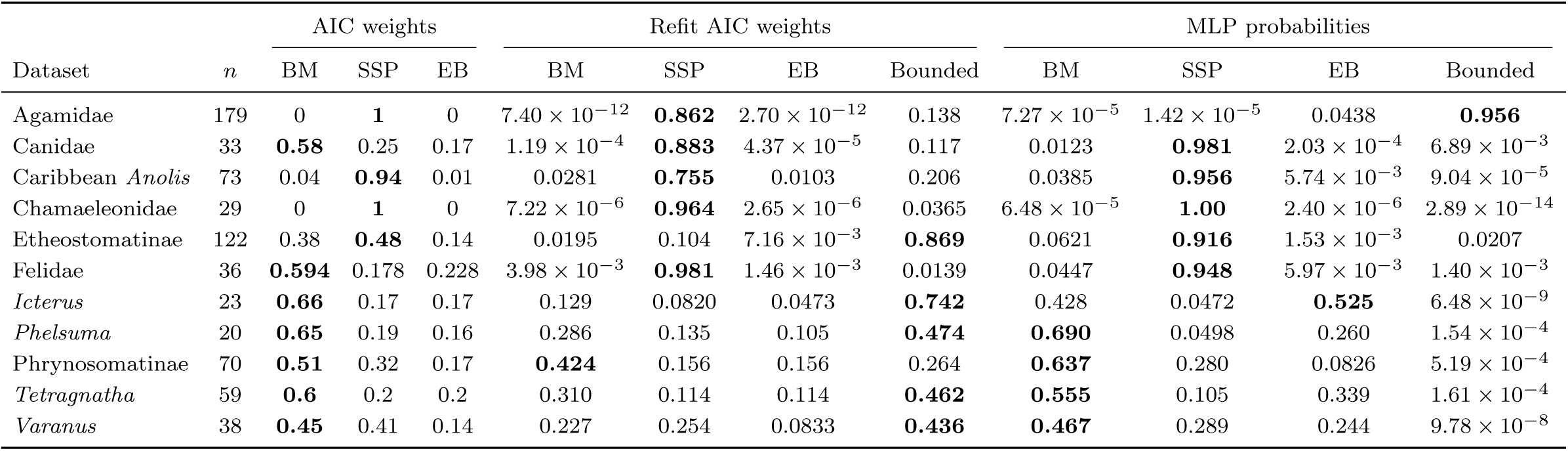
Comparison of model fitting analyses using likelihood-based AIC weights (Harmon et al., 2010b), AIC weights from refitting an expanded candidate set that included bounded BM, and MLP classification probabilities, fitted to quantify **body shape** evolution. AIC weights from Harmon et al. (2010b) compared three models: BM, SSP, and EB. Refit AIC weights and MLP probabilities covered four models: BM, SSP, EB, and bounded BM. *n* is the number of species (tips) in each dataset. For each set of analyses, the dominant model is indicated in bold. All data is taken from the supplement material of Harmon et al. (2010b). **Alt text** Table 5: Model support for body shape evolution in each dataset, shown as likelihood-based AIC weights from Harmon et al. (2010b), AIC weights from refitting an expanded candidate set including bounded BM, and MLP classification probabilities for BM, SSP, EB and bounded BM. The number of species is given for each dataset, and the dominant model in each set of analyses is in bold.

|  |  | AIC weights |  |  | Refit AIC weights |  |  |  | MLP probabilities |  |  |  |
|---|---|---|---|---|---|---|---|---|---|---|---|---|
| Dataset | *n* | BM | SSP | EB | BM | SSP | EB | Bounded | BM | SSP | EB | Bounded |
| Agamidae | 179 | 0 | **1** | 0 | 7.40 × 10<sup>−12</sup> | **0.862** | 2.70 × 10<sup>−12</sup> | 0.138 | 7.27 × 10<sup>−5</sup> | 1.42 × 10<sup>−5</sup> | 0.0438 | **0.956** |
| Canidae | 33 | **0.58** | 0.25 | 0.17 | 1.19 × 10<sup>−4</sup> | **0.883** | 4.37 × 10<sup>−5</sup> | 0.117 | 0.0123 | **0.981** | 2.03 × 10<sup>−4</sup> | 6.89 × 10<sup>−3</sup> |
| Caribbean *Anolis* | 73 | 0.04 | **0.94** | 0.01 | 0.0281 | **0.755** | 0.0103 | 0.206 | 0.0385 | **0.956** | 5.74 × 10<sup>−3</sup> | 9.04 × 10<sup>−5</sup> |
| Chamaeleonidae | 29 | 0 | **1** | 0 | 7.22 × 10<sup>−6</sup> | **0.964** | 2.65 × 10<sup>−6</sup> | 0.0365 | 6.48 × 10<sup>−5</sup> | **1.00** | 2.40 × 10<sup>−6</sup> | 2.89 × 10<sup>−14</sup> |
| Etheostomatinae | 122 | 0.38 | **0.48** | 0.14 | 0.0195 | 0.104 | 7.16 × 10<sup>−3</sup> | **0.869** | 0.0621 | **0.916** | 1.53 × 10<sup>−3</sup> | 0.0207 |
| Felidae | 36 | **0.594** | 0.178 | 0.228 | 3.98 × 10<sup>−3</sup> | **0.981** | 1.46 × 10<sup>−3</sup> | 0.0139 | 0.0447 | **0.948** | 5.97 × 10<sup>−3</sup> | 1.40 × 10<sup>−3</sup> |
| *Icterus* | 23 | **0.66** | 0.17 | 0.17 | 0.129 | 0.0820 | 0.0473 | **0.742** | 0.428 | 0.0472 | **0.525** | 6.48 × 10<sup>−9</sup> |
| *Phelsuma* | 20 | **0.65** | 0.19 | 0.16 | 0.286 | 0.135 | 0.105 | **0.474** | **0.690** | 0.0498 | 0.260 | 1.54 × 10<sup>−4</sup> |
| Phrynosomatinae | 70 | **0.51** | 0.32 | 0.17 | **0.424** | 0.156 | 0.156 | 0.264 | **0.637** | 0.280 | 0.0826 | 5.19 × 10<sup>−4</sup> |
| *Tetragnatha* | 59 | **0.6** | 0.2 | 0.2 | 0.310 | 0.114 | 0.114 | **0.462** | **0.555** | 0.105 | 0.339 | 1.61 × 10<sup>−4</sup> |
| *Varanus* | 38 | **0.45** | 0.41 | 0.14 | 0.227 | 0.254 | 0.0833 | **0.436** | **0.467** | 0.289 | 0.244 | 9.78 × 10<sup>−8</sup> |

The MLP classification altered the distribution of model support, measured as probabilities assigned to each class. Contrary to predictions, the MLP did not reallocate support from BM to EB or from SSP to bounded BM: BM was favored in 14 datasets, SSP in 12, bounded BM in two, and EB in one (Fig. 6, Tables 4 and 5). Although BM was commonly favored, it received relatively low support (maximum *P* (*BM*) = 0.690,*Phelsuma* body shape). Although only a single dataset was assigned to EB under MaxL and MLP classifications, the identity shifted from *Varanus* body size under MaxL to *Icterus* body shape (*P* (EB) = 0.525) under MLP (Table 5). I also hypothesized that the inclusion of bounded BM in the candidate pool may draw support away from SSP/OU1 under MLP. Compared to the original study, this was not common, with only agamid shape evolution showing such a pattern (SSP weight = 1 vs *P* (Bounded) = 0.956). This was one of the few datasets for which addition of bounded BM to the pool under MaxL did not flip support from SSP. In general, the MLP was more conservative in assigning support to bounded BM in comparison to MaxL (2 classifications as best vs 20), which matches expectations based on my simulation study (Fig. 2). Therefore, it appears that EBs are indeed rare, at least in the comparative data compiled by Harmon et al. (2010b).

## Discussion

### A comparison of likelihood-based and MLP evolutionary model classification

Comparative biologists have long used likelihood-based methods to evaluate statistical models that represent distinct evolutionary dynamics despite recognition of their limitations, including the influence of tree size on parameter estimation (Boettiger et al., 2012; Münkemüller et al., 2012; Pennell et al., 2015; Silvestro et al., 2015; Cooper et al., 2016b,a), reduced power to discriminate among models when generating parameters are of low magnitude (Lande, 1976; Hansen, 1997; Butler and King, 2004; Boettiger et al., 2012; Ho and Ané, 2014; Cooper et al., 2016b), and statistical similarity among models representing distinct evolutionary processes (Foote, 1996b; Harmon et al., 2003; Revell et al., 2008; Boucher et al., 2014; Uyeda et al., 2015). The limitations listed above motivate the MLP developed here, which can be evaluated against these biases to determine whether it is a fruitful alternative for evolutionary model classification in comparative studies.

Statistical power depends on sample size for essentially any inferential method that operates in a probabilistic framework, including likelihood ratio tests, information criteria, and Bayesian model comparison (Sokal and Rohlf, 1995; Burnham and Anderson, 2002). Small samples produce flatter likelihood surfaces, greater uncertainty in parameter estimates, and reduced ability to discriminate among candidate models. For example, Boettiger et al. (2012) showed that the reduced capacity to distinguish OU1 from BM on small trees is accompanied by greater uncertainty in phylogenetic signal, with MaxL estimates of Pagel’s *λ* spanning the full parameter space (0 ≤ *λ*^^^ ≤ 1) across simulated datasets on a 13 tip *Geospiza* phylogeny. Because species are phylogenetically non-independent, effective sample size *N*_eff_ ≤ *N* and depends on tree structure; for the 13-species *Geospiza* phylogeny, *N* = 13 is only an upper bound (Ané, 2008) and so tree size is best treated as a proxy for the information available in the data. My bayesian regression results corroborate these effects for MaxL: the relationship between *α* and OU1 classification accuracy differed substantially across tree-size percentiles (Fig. 3A), and shifting from the 10th to 50th percentile increased both maximum model support and the rate at which support increased with *α*. In strong contrast, MLP accuracy showed a much weaker dependence on tree size. It is reasonable to question whether such a comparison is fair, as the MLP takes a fixed-length vector of summary statistics describing the tree and trait data, so every dataset supplies the same number of input values regardless of tree size. This does not circumvent the cause of the sample size problem, however, as smaller trees still contain less information than larger trees and should therefore yield noisier summary statistics. Indeed, my regressions do detect some tree-size dependence in MLP accuracy but, because the MLP learns predictive features directly from the data rather than assuming a multivariate normal distribution and estimating parameters from a flattened likelihood surface, the reduced information in low *N* datasets degrades its performance less severely. This pattern is clear in the posterior predictive plots for MLP performance: the 50th and 90th tree-size percentile curves are largely indistinguishable across generative model classes (Fig. 3D-F). Although the 10th percentile curves are mostly non-overlapping with the others, they remain closer to the higher tree-size percentiles than their corresponding curves under MaxL classification.

The magnitude of model parameters determines how strongly the evolutionary process represented by a given statistical model is expressed in the resulting comparative data and, consequently, the risk of conflation with alternative models. Conflation between BM and alternative quantitative trait models is unavoidable because BM is a special case of all of them. Under OU1 (Lande, 1976; Hansen, 1997), the expected trait distribution and variance-covariance structure converge on those of BM as *α* → 0, flattening the likelihood surface and making parameter optimization difficult (Butler and King, 2004; Ho and Ané, 2014; Cooper et al., 2016b). This nesting has two consequences for likelihood-based inference. First, power declines as parameters approach zero, because data generated under the alternative become nearly indistinguishable from BM (Fig. 3A) and, second, nesting limits how strongly BM can be supported, even when it is the generating process. Because each alternative can reduce to BM, its maximized likelihood can never be lower than that of BM, and when BM generated the data the two will be nearly equal. The difference in AIC is then determined almost entirely by the parsimony penalty, which for OU1, AC, and DC is a single additional parameter (2 AIC units) and for bounded BM is two (4 AIC units) (Boucher and Démery, 2016). BM can therefore be weakly but never strongly preferred over alternative models, relegating it to null model status. The confusion matrix for AIC-based model selection (Fig. 2) shows this clearly, with BM correctly identified in only 34% of cases, and misclassified as OU1 in 6% of cases and bounded BM in a surprising 58% of cases. The MLP, in contrast, correctly identifies a BM model in 78% of cases and never misclassifies it as OU1 or bounded BM (Fig. 2). Again, because the MLP learns the characteristic signature of each model directly from simulated training data (Fig 5), it treats BM as one class among equals rather than as a default null that every alternative can match, and it is not constrained by a parsimony penalty.

Some evolutionary models cannot be distinguished using likelihood-based model evaluation. For example, Uyeda et al. (2015) showed that OU1 and AC yield identical variance-covariance structures on ultrametric trees. The two models are therefore unidentifiable using likelihood-based criteria (Uyeda and Harmon, 2014). Initially, it appeared that these two models were identifiable via MLP (Fig 2). However, fair evaluation of the MLP’s OU1-AC discrimination ability requires data to be subset to the region of parameter space in which likelihood equivalence is expected: *α* ≤ 0.5*a* for true OU1 and σ^2^_0_ ≤ σ^2^_OU1_exp(−*a*) for true AC. Regression of the assigned model probabilities against the corresponding rate parameter for test data within this parameter space revealed near-identical AC–OU1 classification distributions across the two true generating models (Fig. 4) and a bias towards higher *P* (AC) than *P* (OU1). This property cannot be explained by different AC-OU1 ratios in the training data from which the classifier is derived: of the data that fall into the equivalence region, 48.3% are OU1 and 51.7% are AC. Instead, for unknown reasons, the MLP appears to resolve the tie by a tendency to prefer AC over OU1. Unlike MaxL, no parameters are estimated or likelihoods optimized, so this tie is not being broken on the basis of optimizer behavior or parameter bounds. Critically, it is still being broken arbitrarily, albeit in the opposite direction to MaxL where most AC datasets are misclassified as OU1, and in neither case should assignments of non-identifiable models be taken as positive evidence in favor of one or the other process.

A different challenge emerges when attempting to discriminate bounded BM and OU1. Both models produce characteristically low phylogenetic signal patterns in trait data (Foote, 1996b; Revell et al., 2008; Boucher et al., 2014) and are difficult to distinguish in a comparative framework, despite their connections to very different evolutionary processes. Jones and Slater (2026) found asymmetric conflation, where true OU1 was frequently misclassified as bounded BM, particularly for small trees (*n_tip_ <* 100) and a low strength of selection, whereas the reverse of bounded BM being misclassified as OU1 was rare. My results show that MaxL inference of bounded BM also incorrectly absorbs support from true BM, AC, and true DC (Fig. 2), consistent with bounded BM functioning as a “wastebasket” model. This interpretation is further strengthened by the results obtained from adding bounded BM to the candidate pool in the empirical example (Fig. 6). The MLP did not demonstrate a preferential bias toward bounded BM however: the model received no support for true BM, AC, or DC simulations and only 1% support for true OU1 simulations (Fig. 2). This improved discrimination likely reflects the diversity of trait and tree-based summary statistics available to the MLP (Tables 2, 3). Notably, tip excess kurtosis was the second-highest predictor of bounded BM (Fig. 5), consistent with the increasingly uniform trait distributions expected as elapsed time approaches and, ultimately, exceeds the process’ characteristic time and the bounds are realized, whereas BM and OU1 produce approximately Gaussian distributions (Boucher et al., 2014). Drawing on statistics beyond the likelihood therefore helps the MLP in separating models with similar likelihoods and navigating the many-to-one mapping between evolutionary process and phenotypic patterns (Uyeda et al., 2015; Slater and Pennell, 2014; Pennell et al., 2015; Jones and Slater, 2026).

### Empirical case study: detecting signals of early burst dynamics in comparative data

An MLP that is more accurate and less biased than MaxL allows for greater trust in empirical applications and inferred evolutionary processes. Revisiting the claim of Harmon et al. (2010b) that EB events are rarer in comparative data than suggested by evolutionary theory (Simpson, 1944), I asked whether their failure to detect EBs in comparative data reflects a failure of MaxL methods and whether a preference for SSP may reflect a lack of more appropriate, yet difficult to discriminate models, notably bounded BM. Intriguingly, the MLP generated novel class probability distributions that failed to support the first hypothesis and limited support for the second. The EB model was favored only for one dataset based on MLP, though this was a different one than that preferred by MaxL. This result strengthens the conclusion of Harmon et al. (2010b) by showing that their result was not an artifact of the method used.

However, it should be noted that only 29 of the original 88 datasets were of sufficient size (*Ntips* ≥ 20) to evaluate based on the training set provided to my MLP. Similar concerns about clade sizes within this sample, as well as the applicability of EB expectations to small clades or low taxonomic levels have been raised before (Slater and Pennell, 2014; Slater and Friscia, 2019) and the generality of their conclusions to macroevolution remain to be fully evaluated.

In contrast to the dominance of bounded BM support when fitting an expanded suite of models using MaxL, bounded dynamics were preferred for only two datasets under MLP classification (Agamidae body shape and Phrynosomatinae body size). The original AIC weights for the Agamidae were assigned entirely to SSP, whereas the other favored BM. The Agamidae result is consistent with my second hypothesis, that preference for SSP may result from the lack of inclusion of a bounded BM model, though I did not find widespread support for such shifts in these datasets using. The ability to correctly identify a true bounded BM, as well as to reduce the elevated false positive rate that occurs for this model under MaxL, adds further confidence to the classification assigned to this dataset and is a manifestation of the diverse treeand trait-based summary statistics used to train the MLP. Such thorough representations of comparative data are imperative if our interpretations are to reflect the full spectrum of evolutionary dynamics that comparative methods aim to capture and ML approaches provide a meaningful yet computationally cheap way of obtaining and integrating them.

### The future of flexible, mixed-mode evolutionary modeling

Evolutionary patterns arise from interacting processes that operate at different levels of the phylogenetic hierarchy (Jablonski, 2007). The outcomes of these processes are shaped in part by the ecological opportunities available to lineages, but ecological opportunities alone do not determine evolutionary outcomes, and the potential response of any population to novel opportunity is dependent on the developmental and physiological constraints that limit the direction and magnitude of phenotypic change that can be accommodated in response to selection (Gould and Lewontin, 1979; Hansen, 1997; Lande, 1976). Thus, macroevolutionary patterns do not arise in isolation, but rather represent interactions among ecological opportunity, evolutionary constraint, and other sources of evolutionary change (Alencar et al., 2024). Moreover, because different combinations of processes can generate the same pattern (Uyeda et al., 2015; Louca and Pennell, 2020; Jones and Slater, 2026), a clade that experienced constraint in one regime and adaptive differentiation in another may be fit equally well but incorrectly by a single model. Capturing this heterogeneity requires models that represent distinct regimes within a clade and distinguish among their generative processes but existing multi-regime models rarely allow the process itself to vary among regimes. As a result, much of the variation in evolutionary process across the tree of life remains captured. ML classifiers can address this gap because they do not require an explicit likelihood (Breiman, 2001), or a single model across evolutionary regimes. The MLP presented here was trained using single regime univariate models across a broad range of phylogenetic topologies and tree sizes, allowing it to generalize across comparative datasets unlike other ML classifiers such as Evolutionary Discriminant Analysis (EvoDA; Roa Lozano et al., 2026). However, the framework readily extends to multi-regime models, multivariate data and to parameter estimation, either using a regression-based approach or by classifying the model and then optimizing the likelihood. Critically, the MLP approach also extends naturally to mixed-mode, multi-regime inference, in which evolutionary regimes mapped across a phylogeny are classified independently. Graph neural networks (GNNs), particularly graph attention networks (GATs; Kipf and Welling, 2017; Veličković et al., 2018), have further potential for performing this task directly from labeled phylogenies and trait data. Although developing a combined GNN-MLP framework would incur greater computational cost during model training than the single-regime MLP, this cost is incurred only once, and the burden of repeated likelihood optimization across multiple regimes would be alleviated. Extending the present MLP to accommodate heterogeneous evolutionary modes across a phylogeny is therefore an important next step toward scalable inference of multifaceted evolutionary dynamics.

## Conclusion

The MLP framework presented here provides a robust and widely applicable methodological pipeline for evolutionary model classification that substantially alleviates many of the biases associated with alternative MaxL approaches. Empirical application of the MLP supported the scarcity of EB events in comparative data while exemplifying the tendency of MaxL methods to both conflate and favor certain evolutionary models, hindering meaningful evolutionary inference. Subsequent extension of the framework to infer mixed-mode evolutionary scenarios, in which evolutionary modes vary across a phylogeny, will broaden the scope of ML in comparative biology and more faithfully represent the underlying complex macroevolutionary processes. Embracing this complexity will provide a more thorough understanding of how phenotypic traits evolve and how evolutionary dynamics vary within and among clades.

## Supporting information

This document contains the description of the extended methods summarized in the main text, along with supplemental figures (S1) and tables (Table S1-

## Data availability

Scripts and data used to run the analyses are available at the following Dryad repository: Dryad reviewer link

## Acknowledgements

I thank Graham J. Slater for his helpful feedback at various stages of this project. I thank Bailey Howell and the University of Chicago’s Research and Computing Center (RCC) for providing advice and access to computational resources used for data simulations in this study.

