## Supplementary material for "A machine learning framework for evolutionary model classification": This document contains the description of the extended methods summarized in the main text, along with supplemental figures (S1) and tables (Table S1-

Menna Jones

**This document contains the description of the extended methods summarized in the main text, along with supplemental figures (S1) and tables (Table S1-S4).**

### Extended methods

#### K-fold cross validation

After defining the structure of the MLP, architecture and training hyperparameters were optimized using a 10-fold cross-validation procedure. An initial grid search evaluated combinations of learning rate $\in \{5 \times 10^{-4}, 10^{-3}\}$, batch size $\in \{32, 64\}$, weight decay $\in \{0, 10^{-4}\}$, hidden layer size $\in \{32, 64, 128\}$ and dropout rate $\in \{0, 0.15, 0.30\}$, resulting in 96 candidate models. I selected grid parameter values that reflect the overall complexity of the classification task. Hyperparameter combinations were evaluated based on both mean classification accuracy across folds and between-fold variation to ensure both sufficient classification accuracy and precision. Classification accuracy ranged from 74.2-74.6% across the 96 combinations evaluated (Table S1). Between-fold variation was comparably low and standard deviations in accuracy spanned a mere 0.401-0.710%, indicating a network performance that is largely insensitive to moderate changes in training parameters. Because no single hyperparameter combination produced a clear improvement in both predictive accuracy and stability, the final model was selected to balance these two metrics. The final classifier consisted of two hidden layers containing 64 neurons, a learning rate of $5 \times 10^{-4}$, no weight decay, a dropout regularization of $p = 0.15$, and a batch size of 64.

#### Bayesian beta regression

Bayesian beta regression models were constructed to assess issues of model discriminability within MLP and MaxL classification frameworks. For MLP classification, the probability assigned to the generative model was defined as the continuous outcome and regressed onto tree size and generating parameters using Bayesian beta regression with a logit link, fitted via the `brms` package (Bürkner, 2017). Beta regression requires outcomes to fall within the 0-1 interval so probabilities were transformed to $y' = (y(n-1)+0.5)/n$ to avoid boundary values, where $n$ is the sample size (Smithson and Verkuilen, 2006). Thin-plate regression splines were used to accommodate nonlinearity in the marginal effects of each predictor (Wood, 2003). For models with two or more generating parameters (bounded BM, OU1, AC, DC), tensor product smooths were constructed for each parameter-tree size pairing to examine how the influence of each generating parameter on the probability assigned to the true model varied as a function of tree size (Wood, 2003). Weakly informative normal priors ($\mathcal{N}(0, 2)$) were placed on all coefficients, and four MCMC chains of default 2000 iterations (1000 warm-up) were run. Posterior predictive distributions were derived and visualized for all parameter-tree size interaction surfaces whose 95% credible intervals excluded zero. All models converged successfully ($\hat{R} \leq 1.01$, ESS $> 400$ throughout), with minimal divergent transitions. I repeated this approach for the MaxL classification approach with the assigned AICc weight of the true model as the modeled outcome and posterior plots again generated to examine the reliable parameter-tree size interactions.

#### Bayesian Additive Regression Trees (BART)

BART is a nonparametric ensemble ML method that models complex, nonlinear relationships between predictor variables and an outcome variable through a sum of regression trees (Chipman et al., 2010). Each tree recursively partitions the predictor space through binary splits and assigns associated parameter values to terminal nodes. It differs from a traditional random forest algorithm in the placement of prior distributions on all unknown parameters, which function to favor small trees with stable node parameter estimates and ensure that no one tree dominates the model (Bleich et al., 2014).

I constructed a class-specific BART framework to quantify the relative contribution of the twenty summary statistics ($n_{var} = 20$) to MLP classification performance. Using the functions in the `bartMachine` package (Kapelner and Bleich, 2016), separate BART models were trained for each true evolutionary model class, considering the summary statistics as individual predictor variables and the MLP probability assigned to the correct model class as the response variable. Because MLP classification probabilities are bounded between zero and one, values were constrained away from the boundaries ($\epsilon = 10^{-6}$) and logit-transformed prior to BART model fitting. Each dataset was randomly divided into training (80%) and testing (20%) subsets. BART model hyperparameters were chosen using cross-validation across ensemble sizes ($m = 10, 20, 30$), and regularization strength parameters ($k = 1, 2, 3$) using the `bartMachineCV` function. Previous work demonstrates that using a relatively small ensemble size encourages predictors to compete for inclusion in the trees, thereby improving the identification of the most influential variables (Chipman et al., 2010; Bleich et al., 2014). Predictive performance was evaluated using root mean squared error (RMSE) and mean absolute error (MAE) calculated on held-out test data after transforming predictions back to probability space. Final BART models were selected based on the combination of predictive performance and model stability (Table S3). The bottleneck parameter was set to half the total number of trees to ensure adequate representation of the ensemble.

Variable importance was quantified using the `investigate_var_importance`() function, which measures the proportion of times each variable is chosen for a splitting rule across the ensemble of regression trees. Variable rankings were compared across replicate calculations ($n = 20, 50, 100$) to determine the minimum number of replicates needed for convergence. For the final variable importance calculations, ten repetitions were calculated and proportions were averaged across them to account for stochasticity in model fitting. Predictors were ranked according to their mean split frequency and interpreted against the null scenario in which each of the $n_{pred}$ predictors is equally important in splitting rules ($p_{var} = 1/n_{pred} = 0.05$).

Across model classes, BART models recovered non-uniform distributions of predictor importance (Fig. ??, Table S4), with inclusion proportions ranging from from 0.011 for Pagel's $\lambda$ to 0.234 for Blomberg's $K$, both for the DC model. Differences among model classes were evident in both the relative importance and inclusion frequency of individual predictors with no single predictor found to consistently dominate splitting rules. For example, the Mantel statistic was used in only 0.029 of splits for the BM model but 0.087 of DC splitting rules. Similarly, the identity of the most influential tree-based predictor varied among models: the Mantel statistic was the fifth most influential predictor for OU1 and AC, whereas Eigenvector entropy, Rogers' index, and PIC coefficient of variation occupied this position for BM, bounded BM and DC, respectively (Fig. ??). The variable importance distribution for bounded BM model classification is the most unique. Excess kurtosis of the tip-trait distribution and PICs rank as the second- and fourth-most influential predictors, respectively, yet neither appears among the five most influential predictors for BM, OU1, or AC classification. The distribution also appears comparatively uniform and shows only a modest decline in inclusion proportion from the most- to least-influential variables. Several consistent patterns emerged across most or all models. At least three of the five most influential predictors for each model were trait-based, although their identities varied among model classes. For all models except DC, only five predictors exhibited greater influence than expected under the null model of uniform predictor importance, in which each predictor is assumed to contribute independently to the classification outcome.

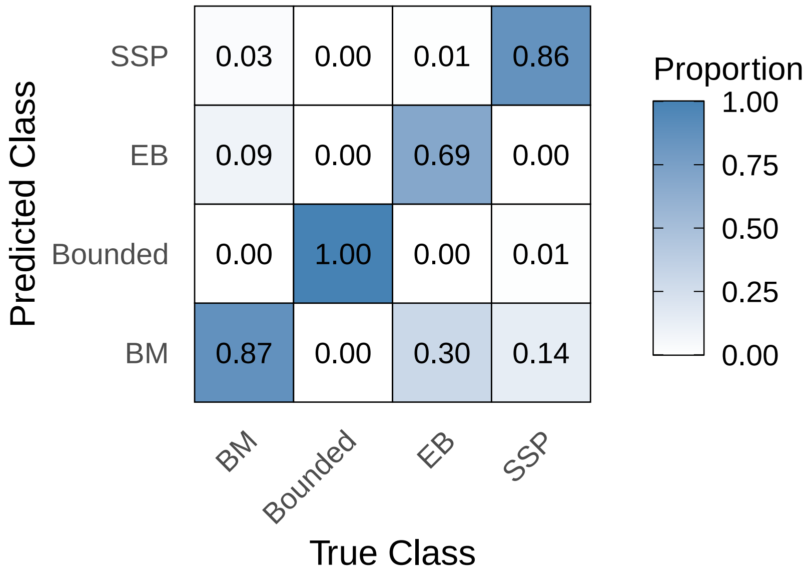


Figure S1: Confusion matrix for the four-class MLP designed for empirical application, evaluated on the test dataset. Rows show the true model class and columns show the predicted class; each row sums to one. To match the empirical analyses, OU1 is labeled SSP and DC is labeled EB.

**Alt text Fig. S1:** Heat map of classification results for the four-class MLP designed for empirical application. Rows are the true model class and columns are the predicted class. Each cell gives the proportion of test cases of that true class assigned to that predicted class. White tiles indicate no classifications and dark blue tiles indicate that (nearly) all cases of that class were assigned there All model classes were correctly classified in the majority of cases..

Table S1: Mean classification accuracy and standard deviation across ten cross-validation folds and 96 candidate MLP models (five-class), defined by unique hyperparameter combinations. Rows are ranked by descending mean accuracy and the final parameter choices are indicated in bold.

| Grid | LR | Weight decay | Dropout | $Hidden_1$ | $Hidden_2$ | Batch | Mean ± SD |
|---|---|---|---|---|---|---|---|
| 93 | $1 \times 10^{-3}$ | 0 | 0.30 | 128 | 64 | 64 | 0.74628 ± 0.00506 |
| 94 | $5 \times 10^{-4}$ | 0 | 0.30 | 128 | 64 | 64 | 0.74624 ± 0.00525 |
| **78** | $5 \times 10^{-4}$ | **0** | **0.15** | **64** | **64** | **64** | **0.74610 ± 0.00465** |
| 70 | $5 \times 10^{-4}$ | 0 | 0.30 | 128 | 32 | 64 | 0.74596 ± 0.00614 |
| 75 | $1 \times 10^{-3}$ | $1 \times 10^{-4}$ | 0.00 | 64 | 64 | 64 | 0.74590 ± 0.00554 |
| 53 | $1 \times 10^{-3}$ | 0 | 0.15 | 64 | 32 | 64 | 0.74589 ± 0.00469 |
| 90 | $5 \times 10^{-4}$ | 0 | 0.15 | 128 | 64 | 64 | 0.74587 ± 0.00656 |
| 46 | $5 \times 10^{-4}$ | 0 | 0.30 | 128 | 64 | 32 | 0.74586 ± 0.00467 |
| 22 | $5 \times 10^{-4}$ | 0 | 0.30 | 128 | 32 | 32 | 0.74584 ± 0.00491 |
| 72 | $5 \times 10^{-4}$ | $1 \times 10^{-4}$ | 0.30 | 128 | 32 | 64 | 0.74571 ± 0.00635 |
| 58 | $5 \times 10^{-4}$ | 0 | 0.30 | 64 | 32 | 64 | 0.74569 ± 0.00490 |
| 68 | $5 \times 10^{-4}$ | $1 \times 10^{-4}$ | 0.15 | 128 | 32 | 64 | 0.74561 ± 0.00516 |
| 56 | $5 \times 10^{-4}$ | $1 \times 10^{-4}$ | 0.15 | 64 | 32 | 64 | 0.74559 ± 0.00545 |
| 96 | $5 \times 10^{-4}$ | $1 \times 10^{-4}$ | 0.30 | 128 | 64 | 64 | 0.74559 ± 0.00585 |
| 80 | $5 \times 10^{-4}$ | $1 \times 10^{-4}$ | 0.15 | 64 | 64 | 64 | 0.74548 ± 0.00495 |
| 16 | $5 \times 10^{-4}$ | $1 \times 10^{-4}$ | 0.00 | 128 | 32 | 32 | 0.74547 ± 0.00554 |
| 51 | $1 \times 10^{-3}$ | $1 \times 10^{-4}$ | 0.00 | 64 | 32 | 64 | 0.74547 ± 0.00588 |
| 54 | $5 \times 10^{-4}$ | 0 | 0.15 | 64 | 32 | 64 | 0.74546 ± 0.00584 |
| 3 | $1 \times 10^{-3}$ | $1 \times 10^{-4}$ | 0.00 | 64 | 32 | 32 | 0.74544 ± 0.00491 |
| 30 | $5 \times 10^{-4}$ | 0 | 0.15 | 64 | 64 | 32 | 0.74541 ± 0.00570 |
| 64 | $5 \times 10^{-4}$ | $1 \times 10^{-4}$ | 0.00 | 128 | 32 | 64 | 0.74541 ± 0.00554 |
| 55 | $1 \times 10^{-3}$ | $1 \times 10^{-4}$ | 0.15 | 64 | 32 | 64 | 0.74540 ± 0.00523 |
| 59 | $1 \times 10^{-3}$ | $1 \times 10^{-4}$ | 0.30 | 64 | 32 | 64 | 0.74536 ± 0.00480 |

| Grid | LR | Weight decay | Dropout | $\text{Hidden}_1$ | $\text{Hidden}_2$ | Batch | Mean $\pm$ SD |
|---|---|---|---|---|---|---|---|
| 19 | $1 \times 10^{-3}$ | $1 \times 10^{-4}$ | 0.15 | 128 | 32 | 32 | $0.74534 \pm 0.00425$ |
| 79 | $1 \times 10^{-3}$ | $1 \times 10^{-4}$ | 0.15 | 64 | 64 | 64 | $0.74531 \pm 0.00535$ |
| 91 | $1 \times 10^{-3}$ | $1 \times 10^{-4}$ | 0.15 | 128 | 64 | 64 | $0.74527 \pm 0.00425$ |
| 52 | $5 \times 10^{-4}$ | $1 \times 10^{-4}$ | 0.00 | 64 | 32 | 64 | $0.74525 \pm 0.00509$ |
| 45 | $1 \times 10^{-3}$ | 0 | 0.30 | 128 | 64 | 32 | $0.74521 \pm 0.00555$ |
| 71 | $1 \times 10^{-3}$ | $1 \times 10^{-4}$ | 0.30 | 128 | 32 | 64 | $0.74519 \pm 0.00520$ |
| 41 | $1 \times 10^{-3}$ | 0 | 0.15 | 128 | 64 | 32 | $0.74518 \pm 0.00523$ |
| 42 | $5 \times 10^{-4}$ | 0 | 0.15 | 128 | 64 | 32 | $0.74517 \pm 0.00425$ |
| 86 | $5 \times 10^{-4}$ | 0 | 0.00 | 128 | 64 | 64 | $0.74517 \pm 0.00483$ |
| 77 | $1 \times 10^{-3}$ | 0 | 0.15 | 64 | 64 | 64 | $0.74512 \pm 0.00556$ |
| 84 | $5 \times 10^{-4}$ | $1 \times 10^{-4}$ | 0.30 | 64 | 64 | 64 | $0.74509 \pm 0.00633$ |
| 28 | $5 \times 10^{-4}$ | $1 \times 10^{-4}$ | 0.00 | 64 | 64 | 32 | $0.74503 \pm 0.00595$ |
| 69 | $1 \times 10^{-3}$ | 0 | 0.30 | 128 | 32 | 64 | $0.74502 \pm 0.00492$ |
| 76 | $5 \times 10^{-4}$ | $1 \times 10^{-4}$ | 0.00 | 64 | 64 | 64 | $0.74501 \pm 0.00577$ |
| 66 | $5 \times 10^{-4}$ | 0 | 0.15 | 128 | 32 | 64 | $0.74496 \pm 0.00486$ |
| 92 | $5 \times 10^{-4}$ | $1 \times 10^{-4}$ | 0.15 | 128 | 64 | 64 | $0.74493 \pm 0.00467$ |
| 47 | $1 \times 10^{-3}$ | $1 \times 10^{-4}$ | 0.30 | 128 | 64 | 32 | $0.74486 \pm 0.00598$ |
| 95 | $1 \times 10^{-3}$ | $1 \times 10^{-4}$ | 0.30 | 128 | 64 | 64 | $0.74480 \pm 0.00554$ |
| 31 | $1 \times 10^{-3}$ | $1 \times 10^{-4}$ | 0.15 | 64 | 64 | 32 | $0.74476 \pm 0.00404$ |
| 1 | $1 \times 10^{-3}$ | 0 | 0.00 | 64 | 32 | 32 | $0.74475 \pm 0.00479$ |
| 65 | $1 \times 10^{-3}$ | 0 | 0.15 | 128 | 32 | 64 | $0.74459 \pm 0.00532$ |
| 21 | $1 \times 10^{-3}$ | 0 | 0.30 | 128 | 32 | 32 | $0.74458 \pm 0.00538$ |
| 60 | $5 \times 10^{-4}$ | $1 \times 10^{-4}$ | 0.30 | 64 | 32 | 64 | $0.74457 \pm 0.00570$ |
| 40 | $5 \times 10^{-4}$ | $1 \times 10^{-4}$ | 0.00 | 128 | 64 | 32 | $0.74451 \pm 0.00513$ |
| 74 | $5 \times 10^{-4}$ | 0 | 0.00 | 64 | 64 | 64 | $0.74448 \pm 0.00561$ |

| Grid | LR | Weight decay | Dropout | $\text{Hidden}_1$ | $\text{Hidden}_2$ | Batch | Mean ± SD |
|---|---|---|---|---|---|---|---|
| 27 | $1 \times 10^{-3}$ | $1 \times 10^{-4}$ | 0.00 | 64 | 64 | 32 | 0.74447 ± 0.00611 |
| 67 | $1 \times 10^{-3}$ | $1 \times 10^{-4}$ | 0.15 | 128 | 32 | 64 | 0.74447 ± 0.00531 |
| 7 | $1 \times 10^{-3}$ | $1 \times 10^{-4}$ | 0.15 | 64 | 32 | 32 | 0.74445 ± 0.00533 |
| 34 | $5 \times 10^{-4}$ | 0 | 0.30 | 64 | 64 | 32 | 0.74444 ± 0.00522 |
| 44 | $5 \times 10^{-4}$ | $1 \times 10^{-4}$ | 0.15 | 128 | 64 | 32 | 0.74444 ± 0.00477 |
| 29 | $1 \times 10^{-3}$ | 0 | 0.15 | 64 | 64 | 32 | 0.74443 ± 0.00411 |
| 11 | $1 \times 10^{-3}$ | $1 \times 10^{-4}$ | 0.30 | 64 | 32 | 32 | 0.74440 ± 0.00707 |
| 17 | $1 \times 10^{-3}$ | 0 | 0.15 | 128 | 32 | 32 | 0.74440 ± 0.00501 |
| 57 | $1 \times 10^{-3}$ | 0 | 0.30 | 64 | 32 | 64 | 0.74432 ± 0.00600 |
| 49 | $1 \times 10^{-3}$ | 0 | 0.00 | 64 | 32 | 64 | 0.74429 ± 0.00525 |
| 62 | $5 \times 10^{-4}$ | 0 | 0.00 | 128 | 32 | 64 | 0.74418 ± 0.00462 |
| 61 | $1 \times 10^{-3}$ | 0 | 0.00 | 128 | 32 | 64 | 0.74416 ± 0.00472 |
| 25 | $1 \times 10^{-3}$ | 0 | 0.00 | 64 | 64 | 32 | 0.74415 ± 0.00556 |
| 89 | $1 \times 10^{-3}$ | 0 | 0.15 | 128 | 64 | 64 | 0.74409 ± 0.00534 |
| 85 | $1 \times 10^{-3}$ | 0 | 0.00 | 128 | 64 | 64 | 0.74406 ± 0.00508 |
| 38 | $5 \times 10^{-4}$ | 0 | 0.00 | 128 | 64 | 32 | 0.74393 ± 0.00562 |
| 88 | $5 \times 10^{-4}$ | $1 \times 10^{-4}$ | 0.00 | 128 | 64 | 64 | 0.74390 ± 0.00579 |
| 43 | $1 \times 10^{-3}$ | $1 \times 10^{-4}$ | 0.15 | 128 | 64 | 32 | 0.74387 ± 0.00601 |
| 83 | $1 \times 10^{-3}$ | $1 \times 10^{-4}$ | 0.30 | 64 | 64 | 64 | 0.74386 ± 0.00622 |
| 33 | $1 \times 10^{-3}$ | 0 | 0.30 | 64 | 64 | 32 | 0.74385 ± 0.00573 |
| 63 | $1 \times 10^{-3}$ | $1 \times 10^{-4}$ | 0.00 | 128 | 32 | 64 | 0.74378 ± 0.00564 |
| 81 | $1 \times 10^{-3}$ | 0 | 0.30 | 64 | 64 | 64 | 0.74376 ± 0.00573 |
| 13 | $1 \times 10^{-3}$ | 0 | 0.00 | 128 | 32 | 32 | 0.74371 ± 0.00469 |
| 82 | $5 \times 10^{-4}$ | 0 | 0.30 | 64 | 64 | 64 | 0.74364 ± 0.00617 |
| 35 | $1 \times 10^{-3}$ | $1 \times 10^{-4}$ | 0.30 | 64 | 64 | 32 | 0.74362 ± 0.00476 |

| Grid | LR | Weight decay | Dropout | $\text{Hidden}_1$ | $\text{Hidden}_2$ | Batch | Mean ± SD |
|---|---|---|---|---|---|---|---|
| 37 | $1 \times 10^{-3}$ | 0 | 0.00 | 128 | 64 | 32 | 0.74351 ± 0.00496 |
| 50 | $5 \times 10^{-4}$ | 0 | 0.00 | 64 | 32 | 64 | 0.74349 ± 0.00608 |
| 14 | $5 \times 10^{-4}$ | 0 | 0.00 | 128 | 32 | 32 | 0.74347 ± 0.00560 |
| 87 | $1 \times 10^{-3}$ | $1 \times 10^{-4}$ | 0.00 | 128 | 64 | 64 | 0.74319 ± 0.00590 |
| 15 | $1 \times 10^{-3}$ | $1 \times 10^{-4}$ | 0.00 | 128 | 32 | 32 | 0.74315 ± 0.00665 |
| 73 | $1 \times 10^{-3}$ | 0 | 0.00 | 64 | 64 | 64 | 0.74308 ± 0.00508 |
| 23 | $1 \times 10^{-3}$ | $1 \times 10^{-4}$ | 0.30 | 128 | 32 | 32 | 0.74307 ± 0.00687 |
| 39 | $1 \times 10^{-3}$ | $1 \times 10^{-4}$ | 0.00 | 128 | 64 | 32 | 0.74219 ± 0.00649 |

**Alt text Table S1:** Summary of the 10-fold cross-validation used to define the five-class MLP architecture. 96 models were constructed across combinations of learning rate (LR), dropout, size of hidden layers one and two, and batch size. Model 78 was chosen as the final model due to its high mean accuracy and narrow standard deviation.

Table S2: Bayesian beta regression (logit link) results across true generative models and classification frameworks. For MaxL, the response outcome is the Akaike weight assigned to the true model. For the MLP, the response variable is the probability assigned to the true model. Posterior means and 95% credible intervals are displayed on the logit scale for the intercept, precision parameter $\phi$, and tensor-product ($t_2$) components credible under at least one classifier; estimates whose intervals exclude zero are in bold. Subscripts index components of the tensor-product basis. Components of $t_2(\text{ntip}, \cdot)_2$ in models with two smooths are not separately identified and are omitted. Baseline accuracy is defined as the predicted response at mean covariate values.

| | | **MLP** | | **MaxL** | |
|---|---|---|---|---|---|
| **Model** | **Term** | Estimate | 95% CI | Estimate | 95% CI |
| BM | Intercept | **-0.12** | [-0.16, -0.08] | **-1.29** | [-1.34, -1.24] |
| | Baseline | 0.47 | — | 0.22 | — |
| | $t_2(\text{ntip}, \sigma^2)_1$ | **-0.04** | [-0.07, -0.01] | -0.02 | [-0.07, 0.02] |
| | $t_2(\text{ntip}, \sigma^2)_2$ | **0.26** | [0.23, 0.29] | **0.18** | [0.14, 0.22] |
| | $\phi$ | 10.52 | [9.89, 11.16] | 5.98 | [5.61, 6.36] |
| Bounded BM | Intercept | **5.50** | [5.39, 5.60] | **9.29** | [9.23, 9.34] |
| | Baseline | 0.996 | — | >0.9999 | — |
| | $t_2(\text{ntip}, \tau)_1$ | 0.02 | [-0.03, 0.07] | **0.05** | [0.01, 0.09] |
| | $t_2(\text{ntip}, \tau)_3$ | 0.03 | [-0.03, 0.09] | **0.08** | [0.02, 0.13] |
| | $t_2(\text{ntip}, \sigma^2)_3$ | 0.00 | [-0.06, 0.06] | -0.06$^{\dagger}$ | [-0.12, -0.00] |
| | $\phi$ | 44.58 | [39.76, 49.49] | 14033.48 | [13065.17, 15060.92] |
| OU1 | Intercept | **1.01** | [0.94, 1.08] | **1.18** | [1.11, 1.26] |
| | Baseline | 0.73 | — | 0.77 | — |
| | $t_2(\text{ntip}, \alpha)_1$ | **-1.46** | [-1.53, -1.39] | **1.01** | [0.94, 1.09] |
| | $t_2(\text{ntip}, \alpha)_3$ | **-0.27** | [-0.34, -0.20] | **-0.17** | [-0.25, -0.09] |
| | $\phi$ | 4.20 | [3.91, 4.48] | 1.89 | [1.75, 2.04] |
| AC | Intercept | **-0.19** | [-0.24, -0.15] | **-3.31** | [-3.38, -3.24] |
| | Baseline | 0.45 | — | 0.035 | — |
| | $t_2(\text{ntip}, a)_1$ | **0.33** | [0.29, 0.38] | **0.60** | [0.55, 0.66] |
| | $t_2(\text{ntip}, a)_3$ | **-0.08** | [-0.13, -0.03] | **-0.21** | [-0.27, -0.14] |
| | $\phi$ | 8.33 | [7.83, 8.81] | 17.65 | [16.38, 18.95] |
| DC | Intercept | **0.78** | [0.71, 0.85] | **-1.07** | [-1.11, -1.02] |
| | Baseline | 0.69 | — | 0.26 | — |
| | $t_2(\text{ntip}, a)_1$ | **1.13** | [1.06, 1.20] | **-0.57** | [-0.62, -0.52] |
| | $t_2(\text{ntip}, a)_3$ | **-0.15** | [-0.22, -0.08] | -0.01 | [-0.07, 0.04] |
| | $t_2(\text{ntip}, \sigma^2)_3$ | 0.07$^{\dagger}$ | [0.00, 0.13] | 0.05$^{\dagger}$ | [0.00, 0.10] |
| | $\phi$ | 2.62 | [2.47, 2.78] | 8.53 | [8.04, 9.04] |

**Alt text Table S2:** Bayesian beta regression results for the MLP and MaxL classifiers across true generative models. Columns give the posterior mean estimate and 95% credible interval for the MLP and MaxL classification, measured as probability and Akaike weight assigned to the true model, respectively. For each model, the table displays the intercept, baseline predicted accuracy, credible tensor-product components, and the precision parameter $\phi$. Estimates whose intervals do not span zero are indicated in bold.

Table S3: BART model parameterization chosen through cross-validation: number of trees in the ensemble, *mu*; shrinkage parameter, *k*; degrees of freedom for the prior on the error variance, $\nu$; quantile used to set the prior on the error variance, *q*.

| | $m$ | $k$ | $\nu$ | $q$ |
|---|---|---|---|---|
| **BM** | 20 | 3 | 3 | 0.99 |
| **Bounded BM** | 30 | 3 | 3 | 0.9 |
| **OU1** | 30 | 3 | 3 | 0.9 |
| **AC** | 30 | 3 | 3 | 0.9 |
| **DC** | 30 | 3 | 10 | 0.75 |

**Alt text Table S3:** Final BART model parameterizations used to evaluate variable importance in MLP classification.

Table S4: Variable inclusion proportions from the BART analysis. Top five variables are highlighted in bold for each model. Variable abbreviations: *pagdelta*, Pagel's $\delta$; *blomK*, Blomberg's $K$; *pagkappa*, Pagel's $\kappa$; *pic_cv*, PIC coefficient of variation; *entropy*, eigenvalue entropy; *picexckurt*, PIC excess kurtosis; *vpd*, variation in pairwise distances; *ntip*, number of tips; *rogers*, Rogers' index; *totpath*, total path length; *totcoph*, total cophenetic distance; *mantel*, Mantel statistic; *maxdepth*, maximum tree depth; *paglambda*, Pagel's $\lambda$; *coll*, Colless index; *varbl*, variation in branch length; *picskew*, PIC skewness; *tipskew*, tip skewness; *tipexckurt*, tip excess kurtosis; and *gamma*, $\gamma$ statistic.

| | BM | | Bounded BM | | OU1 | | AC | | DC | |
|---|---|---|---|---|---|---|---|---|---|---|
| Variable | Mean | SD | Mean | SD | Mean | SD | Mean | SD | Mean | SD |
| pagdelta | **0.223** | 0.003 | 0.023 | 0.001 | 0.021 | 0.002 | **0.173** | 0.002 | **0.168** | 0.003 |
| blomK | **0.125** | 0.002 | 0.052 | 0.001 | **0.151** | 0.003 | **0.167** | 0.003 | **0.234** | 0.003 |
| pagkappa | **0.102** | 0.002 | **0.061** | 0.002 | **0.155** | 0.003 | **0.095** | 0.003 | 0.044 | 0.002 |
| pic_cv | **0.072** | 0.002 | 0.053 | 0.001 | **0.090** | 0.002 | 0.054 | 0.001 | **0.056** | 0.002 |
| entropy | **0.060** | 0.002 | 0.042 | 0.001 | 0.035 | 0.001 | **0.098** | 0.002 | 0.030 | 0.002 |
| picexckurt | 0.049 | 0.002 | **0.059** | 0.002 | 0.032 | 0.001 | 0.027 | 0.001 | **0.079** | 0.003 |
| vpd | 0.042 | 0.002 | 0.043 | 0.001 | 0.034 | 0.001 | 0.043 | 0.003 | 0.033 | 0.001 |
| ntip | 0.034 | 0.001 | 0.049 | 0.001 | 0.052 | 0.003 | 0.033 | 0.001 | 0.026 | 0.001 |
| rogers | 0.032 | 0.001 | **0.057** | 0.001 | 0.048 | 0.002 | 0.027 | 0.001 | 0.020 | 0.001 |
| totpath | 0.030 | 0.001 | 0.048 | 0.001 | 0.045 | 0.002 | 0.029 | 0.001 | 0.024 | 0.001 |
| totcoph | 0.029 | 0.001 | 0.051 | 0.001 | 0.041 | 0.002 | 0.024 | 0.002 | 0.022 | 0.001 |
| mantel | 0.029 | 0.001 | 0.039 | 0.001 | **0.059** | 0.002 | **0.057** | 0.002 | **0.087** | 0.002 |
| maxdepth | 0.028 | 0.001 | 0.049 | 0.001 | 0.027 | 0.002 | 0.030 | 0.001 | 0.035 | 0.001 |
| paglambda | 0.027 | 0.001 | 0.040 | 0.001 | 0.028 | 0.003 | 0.019 | 0.001 | 0.011 | 0.001 |
| coll | 0.025 | 0.000 | 0.050 | 0.001 | 0.031 | 0.001 | 0.025 | 0.001 | 0.025 | 0.001 |
| varbl | 0.025 | 0.001 | **0.080** | 0.001 | **0.067** | 0.003 | 0.031 | 0.001 | 0.023 | 0.001 |
| picskew | 0.021 | 0.001 | 0.045 | 0.001 | 0.015 | 0.001 | 0.014 | 0.001 | 0.017 | 0.001 |
| tipskew | 0.019 | 0.001 | 0.046 | 0.001 | 0.016 | 0.001 | 0.017 | 0.001 | 0.032 | 0.001 |
| tipexckurt | 0.016 | 0.001 | **0.077** | 0.001 | 0.039 | 0.001 | 0.012 | 0.001 | 0.022 | 0.001 |
| gamma | 0.013 | 0.001 | 0.037 | 0.001 | 0.014 | 0.001 | 0.023 | 0.001 | 0.015 | 0.001 |

**Alt text Table S4:** Full set of variable inclusion proportions across MLP predictions, partitioned by true generative model. Bold values represent the five variables with the highest inclusion proportion for each model class.
